# Disease-causing mutations in Synaptotagmin can act via dominant-negative, gain-of-function or haploinsufficient mechanisms

**DOI:** 10.64898/2026.03.17.712223

**Authors:** Zhuo Guan, Maria Bykhovskaia, J. Troy Littleton

**Affiliations:** The Picower Institute for Learning and Memory, Department of Brain and Cognitive Sciences, Department of Biology, Massachusetts Institute of Technology, Cambridge, MA 02139; Department of Neurology, Wayne State University, Detroit, MI, 48201

**Author notes:** Correspondence (J.T.L.).

## Abstract

Members of the Synaptotagmin (SYT) family of synaptic vesicle (SV) proteins contain two Ca^2+^ binding C2 domains (C2A and C2B) that regulate the timing and probability of SV fusion. Recently, dominant mutations in SYT1, present at most CNS synapses, and SYT2, abundant in the PNS, have been found to cause human neurological diseases. Most pathogenic alleles localize to the C2B Ca^2+^ binding pocket, although mutations outside of this region have been identified. Here we report the effects of a large array of disease-associated human SYT point mutations on synaptic transmission using the *Drosophila* model to define their mechanism of action. Mutations in the C2B Ca^2+^ binding pocket were the most severe and acted in a dominant-negative manner to decrease evoked release. Indeed, mutations in the C2B Ca^2+^ binding pocket were a privileged site for dominant-negative effects on SV fusion. Molecular dynamics simulations indicate these mutations alter the topology of the C2B Ca^2+^ binding pocket and disrupt Ca^2+^-triggered membrane insertion. Several mutations outside of this region acted through a gain-of-function mechanism to enhance neurotransmitter release, while others caused the protein to undergo degradation. Consistent with this latter subset acting via haploinsufficiency, heterozygotes lacking one copy of SYT1 displayed a ∼40% decrease in evoked release. These data indicate C2B Ca^2+^ binding pocket mutations act dominantly to poison the fusion machinery and block release sites, while mutations outside of this region cause haploinsufficiency or gain-of-function effects with milder effects on synaptic output and behavioral phenotypes.

## Introduction

Synaptic vesicles (SVs) fuse with the plasma membrane (PM) using a highly conserved machinery that includes the SNARE complex and associated regulatory proteins^1,2^. Several members of the Synaptotagmin (SYT) family localize to SVs, including SYT1 and SYT2, and contain two Ca^2+^ binding C2 domains (C2A and C2B) that regulate the timing and probability of SV release^3–6^. Although SYTs have several functions during the SV cycle, their ability to bind SNARE complexes and the PM in a Ca^2+^-independent manner facilitates SV priming^7–10^. Together with the SNARE-binding protein Complexin (CPX), Ca^2+^-independent SYT binding to SNARE complexes also reduces spontaneous fusion by regulating the timing of full SNARE assembly^7,11,12^. These interactions position SYT at the site of membrane fusion and allow the hydrophobic loops that emerge from the C2 domains to bind Ca^2+^ and penetrate into the PM upon Ca^2+^ entry^10,12–20^. Ca^2+^-dependent lipid insertion creates a rotation in SYT conformation that is hypothesized to activate the final stages of SNARE assembly and drive opening of the fusion pore^21–24^. Single vesicle imaging and proteomic approaches indicate 7-20 SYT1 proteins are found on an individual SV^25–27^. Together with the requirement for multiple SNARE complexes to assemble per fusion event^28–30^, current models indicate multiple SYT proteins on a vesicle interact with assembling SNAREs in a highly organized arrangement at the SV-PM interface^6,31^.

Given their key role in synaptic transmission as Ca^2+^ sensors, disruption in SYTs would be predicted to result in neurological phenotypes in humans. The first mutations in a human SYT protein were identified in patients presenting with an autosomal dominant congenital myasthenic syndrome (CMS) that caused peripheral muscle weakness^32^. These patient families carried point mutations in one allele of SYT2, which is abundantly expressed in motoneurons^4^, and had defective neuromuscular junction (NMJ) function but no cognitive defects. More recently, mutations disrupting one allele of SYT1 were identified that caused a more debilitating neurodevelopmental disorder termed Baker-Gordon Syndrome (also known as SYT1-associated neurodevelopmental disorder), with patients showing cognitive dysfunction and developmental delay, often with a failure to develop language^33–39^. These dominant SYT1 mutations are not heritable due to severe cognitive dysfunction, but arise as spontaneous mutations in the germline. Most disease-causing mutations in SYT1 and SYT2 localize to the C2B Ca^2+^ binding pocket and are hypothesized to act as dominant-negative alleles that reduce neurotransmitter release^7,32,36,38,40^. Indeed, disrupting either SNARE binding or Ca^2+^-independent membrane interactions prevent the toxicity of C2B Ca^2+^ binding mutations^7^, indicating they block fusion downstream of SV docking. More recently, mutations outside of the SYT1 C2B Ca^2+^ binding pocket that cause less severe cognitive defects have been identified^41^, raising the possibility of additional pathogenic mechanisms. The challenge from a clinical perspective is to understand how SYT mutations disrupt the fusion machinery and identify ways to normalize neurotransmitter release.

To characterize how SYT1 and SYT2 alleles alter synaptic transmission, we assayed the effects of a large array of disease-associated SYT mutations in *Drosophila*. While both SYT1 and SYT2 localize to SVs in mammals^3^, *Drosophila* contains a single member of the SV subfamily (SYT1)^42^. We found that heterozygotes lacking one copy of SYT1 display a 40% decrease in evoked release, suggesting haploinsufficiency may account for milder phenotypes associated with alleles that cause SYT1 degradation or loss-of-function. In contrast, several mutations in the C2A and C2B domain acted through a gain-of-function mechanism to increase neurotransmitter release. The most severe disease-causing mutations were in the C2B Ca^2+^ binding pocket and dramatically decreased evoked SV fusion. These data indicate C2B Ca^2+^ binding pocket mutants act dominantly to poison SV fusion by allowing SYT1 to engage SNAREs while preventing normal Ca^2+^ activation of the fusion machinery, resulting in a blockage of synaptic release sites.

## Results

### Haploinsufficiency of *Synaptotagmin 1* (*Syt1*) decreases evoked neurotransmitter release

More than 50 heterozygous mutations in SYT1 and SYT2 with proposed clinical relevance have been deposited in NIH’s ClinVar archive, with many more alleles of unknown significance. Although most patients have not been phenotypically described in detail, some have more extensive clinical evaluations^32–37,40,43^. We focused on examining this latter subset to examine how their unique variants in SYT alter synaptic transmission at *Drosophila* NMJs, a well characterized glutamatergic synapse previously used to model synaptic diseases, including those associated with SYT2^32,40,44^. For SYT1, three broad allele categories could be identified based on location of the altered amino acid. These included C2A mutations located outside of its Ca^2+^ binding pocket (L159R, T196K, E209K, E219Q) and C2B mutations located both within (D304G, D366E, I368T) and outside (N341S) of its Ca^2+^ binding pocket (Figure 1A-C). For SYT2, all dominant disease alleles map to the C2B Ca^2+^ binding pocket, including D307A and I371K that are characterized in this study (Figure 1A, D). We compared these alleles to a series of non-disease associated mutations that alter other SYT1 interactions (Figure 1B, E), including those disrupting the primary SNARE-binding interface (S280L), the tripartite SNARE-CPX interface (L388Q, L395Q), the C2B polybasic Ca^2+^-independent lipid binding domain (K327E), C2A-C2B domain interactions (R200H), and the C2A Ca^2+^ binding pocket (D233N). We also compared disease alleles to mutations that conserve or alter the charge or hydrophobicity of the C2A (D233E, D233R, F235E) or C2B (D366N, D366E, D366R) Ca^2+^ binding pockets. For clarity, the human SYT1 amino acid number is used throughout except for SYT2 disease alleles, in which case the human SYT2 amino acid number is used. The corresponding *Drosophila* residue is shown in Figure 1A.

**Figure 1.**
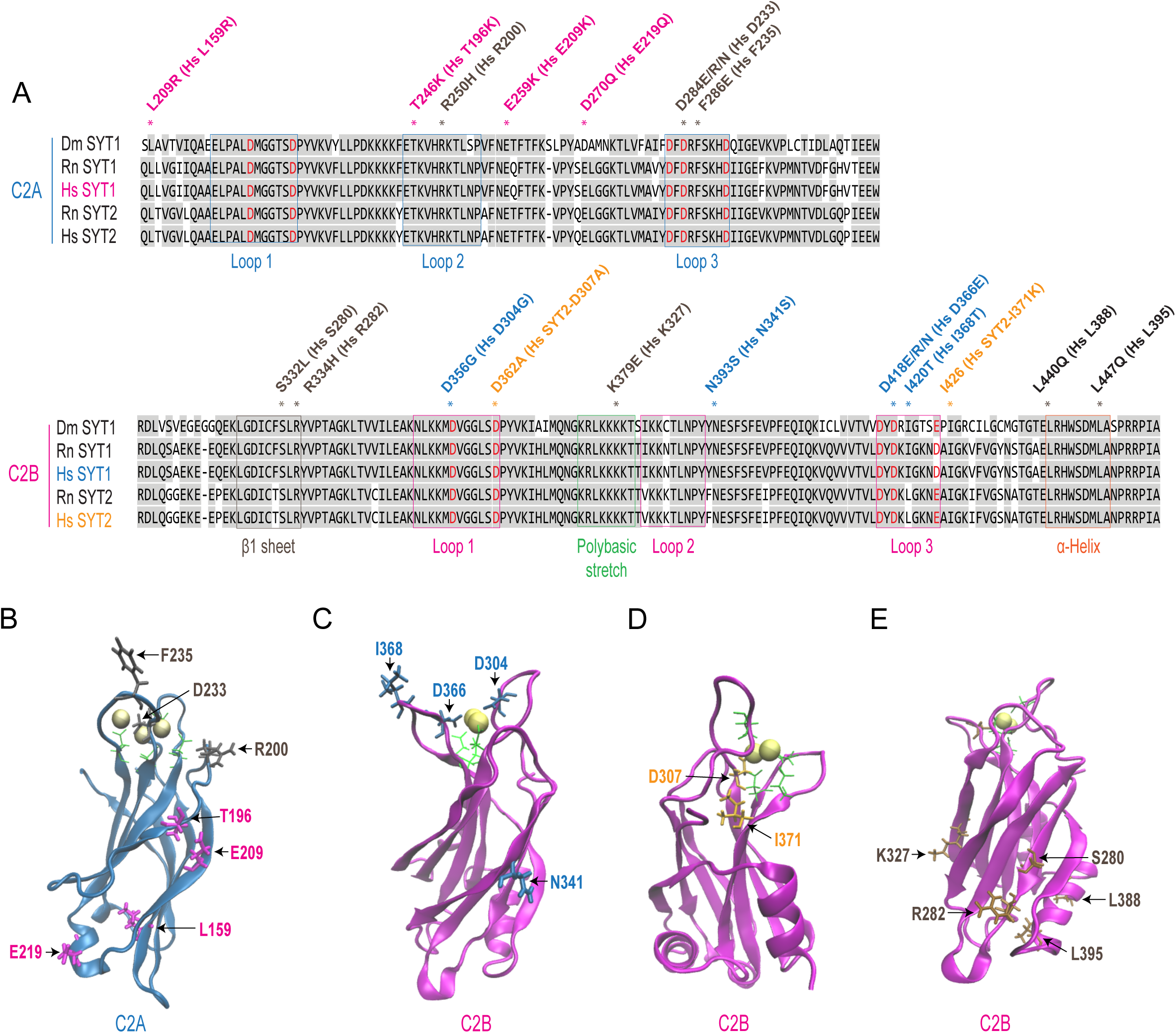
Location of disease associated point mutations in SYT1 and SYT2. (A) Sequence alignment of the C2A and C2B domains of human (Hs) and rat (Rn) SYT1 and SYT2 with *Drosophila* (Dm) SYT1. Conserved resides are highlighted in gray. C2A and C2B Ca^2+^ binding residues are noted in red, with the membrane penetrating loops boxed in blue (C2A) or magenta (C2B). The primary SNARE complex binding interface in the C2B β1 sheet is boxed in grey, the C2B polybasic stretch in green and the C2B tripartite SNARE-CPX binding α-helix in orange. The location of residues examined in this study are denoted with asterisks, including human C2A SYT1 alleles (magenta), human SYT1 C2B alleles (blue), human SYT2 C2B alleles (orange), and mutations disrupting specific SYT1-mediated interactions (black). The *Drosophila* amino acid number is indicated, with the homologous human SYT1 or SYT2 residue in parentheses. (B-E) Structures of the C2A (B) and C2B (C-E) domains of *Drosophila* SYT1 generated with VMD 1.9.3 that show the location of mutated residues indicated with their corresponding human amino acid number. For C2A (B), the structures depict location of human SYT1 alleles and non-disease interaction-specific mutations used in the study. For C2B, the structures depict location of human SYT1 alleles (C), human SYT2 alleles (D) and non-disease interaction-specific mutations (E) used in the study. Ca^2+^ ions are denoted in yellow and the five Ca^2+^ binding aspartate residues are highlighted in green (or blue/orange if representing a mutant allele).

Although C2B Ca^2+^ binding pocket mutations in SYT1 and SYT2 are hypothesized to act in a dominant-negative manner, the identification of SYT1 mutations outside this region suggest the possibility of additional mechanisms. This group of alleles generally result in milder neurodevelopmental phenotypes such as autism, hyperactivity and mild intellectual disability, compared with the C2B Ca^2+^ pocket mutations that are more severe and typically prevent language acquisition^34–36^. Haploinsufficiency, associated with 50% loss of a protein’s function, has been demonstrated for multiple proteins involved in neurotransmission that cause neurodevelopmental disorders, including components of the SNARE machinery^45,46^. These findings raise the possibility that some SYT1 disease alleles may act in a similar fashion. To assay if neurotransmission is sensitive to reduced SYT1 protein levels, we compared release between controls and heterozygotes containing one copy of a *Syt1* null mutation (*Syt1^AD4/+^*) using two-electrode voltage clamp recordings from 3^rd^ instar larval NMJs. *Drosophila* lacking one copy of SYT1 showed a 40% decrease in protein levels by western analysis (Figure 2A, B) and a 44% decrease in SYT1 immunostaining within presynaptic terminals (Figure 2C, D). Although synaptic morphology and NMJ growth were not altered in *Syt1^AD4/+^* heterozygotes (Figure 2E), evoked neurotransmitter release was reduced across a range of extracellular [Ca^2+^] (Figure 2F-H). In 1.5 mM Ca^2+^, the amplitude of evoked responses was reduced by ∼40% in *Syt1^AD4/+^*compared to controls (control: 212.5 ± 7.7 nA; *Syt1^AD4/+^*: 133.7 ± 11.4). Although evoked release was reduced, the Ca^2+^ cooperativity of release (control: 2.76 ± 0.13; *Syt1^AD4/+^*: 2.87 ± 0.08) and the rate of spontaneous miniature excitatory junctional currents (mEJCs) were not altered (control: 1.77 ± 0.16 Hz; *Syt1^AD4/+^*: 2.19 ± 0.26, Figure 2I). We conclude haploinsufficiency for SYT1 decreases evoked neurotransmission, raising the possibility some disease alleles act via this mechanism (see below).

**Figure 2.**
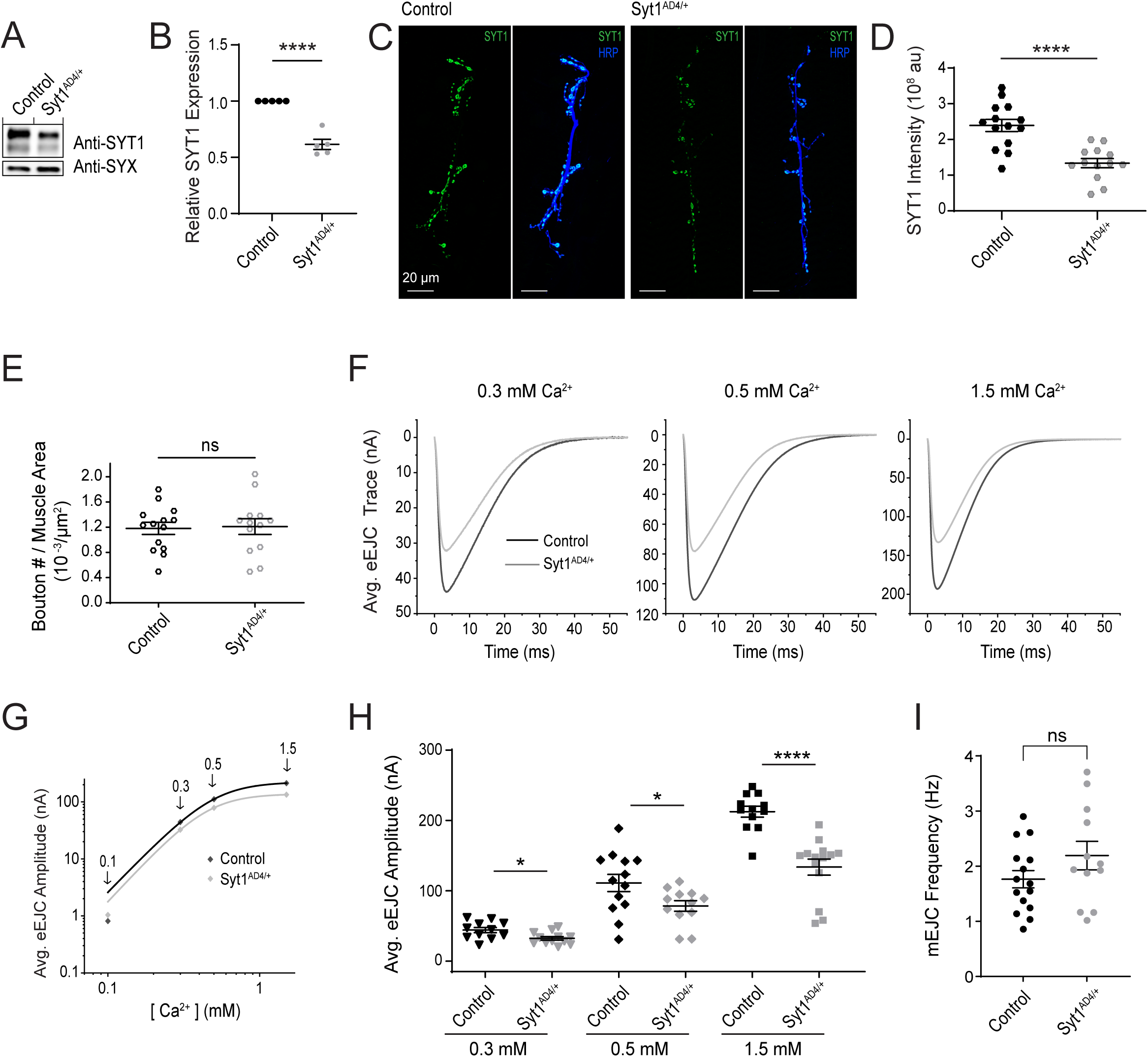
Haploinsufficiency of SYT1 reduces its expression at synapses and decreases evoked release. (A) Representative western of SYT1 protein levels from adult brain extracts of *Syt1* null heterozygotes (*Syt1^AD4/+^*) versus controls. Anti-Syntaxin1 (SYX) was used as a loading control. (B) Quantification of relative SYT1 protein levels in heterozygotes (*Syt1^AD4/+^*) normalized to control. (C) Immunostaining for SYT1 (green) and HRP (blue) at 3^rd^ instar muscle 6/7 NMJs for the indicated genotypes. Note the reduced levels of synaptic SYT1 in the heterozygote. (D) Quantification of total SYT1 fluorescence within the HRP-positive NMJ area (au = arbitrary units). (E) Quantification of varicosity number normalized to muscle surface area for the indicated genotypes. (F) Average eEJC traces recorded from control (black) or *Syt1^AD4/+^* (gray) at the indicated extracellular [Ca^2+^]. (G) Ca^2+^ cooperativity of release is displayed on a double logarithmic plot across the indicated extracellular Ca^2+^ range. Ca^2+^ cooperativity is not affected in *Syt1* heterozygotes, though evoked release is decreased across all [Ca^2+^] levels. (H) Quantification of mean eEJC amplitude at the indicated extracellular [Ca^2+^]. (I) Quantification of mean spontaneous mEJC rate. Mini frequency is not altered by the loss of one *Syt1* allele. Data are shown as mean ± SEM. Statistical significance: \**p* < 0.05, \*\*\*\**p* < 0.0001, ns = not significant. Raw values, statistical tests and sample number are provided in the Source Data file.

### Mutations in the C2B Ca^2+^ binding loops dominantly disrupt evoked release and fail to rescue *Syt1* null phenotypes

Given mutations in the C2B Ca^2+^ binding pocket are proposed to act in a dominant fashion to suppress neurotransmitter release, we hypothesized these alleles would be sensitive to the endogenous levels of wildtype SYT1. As multiple SYT1 proteins are required to drive fusion of a single SV, having more wildtype protein should compete with the mutant version for triggering SNARE-mediated fusion. SYT1’s C2 domains each contain five negatively charged aspartic acid residues (numbered D1 to D5) embedded within two loops that form the Ca^2+^-binding pocket and are critical for Ca^2+^ binding and subsequent membrane penetration. To test if synaptic transmission was sensitive to the balance between wildtype and mutant SYT1, we overexpressed a previously described C2B Ca^2+^ binding mutant where the first and second aspartic acid residues are mutated to asparagine^7,18^. SYT1 protein expression detected by westerns with anti-SYT1 antisera was similar following expression of wildtype UAS-SYT1 or UAS-SYT1^C2B-D304N,D310N^ (hereafter referred to as SYT1^C2BD1,2N^) transgenes (Supplemental Figure 1A, B). Pan-neuronal expression of SYT1^C2BD1,2N^ with *elav*^C155^ GAL4 in an otherwise wildtype *Drosophila* background with two endogenous copies of native SYT1 resulted in a 62% decrease in evoked release. Expression of the same transgene in *Syt1^AD4/+^* heterozygotes caused a more severe 77% decrease in release (Figure 3A, B). Pan-neuronal overexpression of wildtype SYT1 did not reduce release in controls or in *Syt1^AD4/+^*heterozygotes (Figure 3B). SYT1C2B^D1,2N^ also resulted in an increase in spontaneous release only when overexpressed in *Syt1^AD4/+^*heterozygotes (Figure 3C). We conclude the ability of the SYT1 C2B Ca^2+^ binding mutant to decrease evoked release and alter clamping of spontaneous fusion is dosage sensitive and enhanced when endogenous SYT1 levels are reduced in heterozygote conditions.

**Figure 3.**
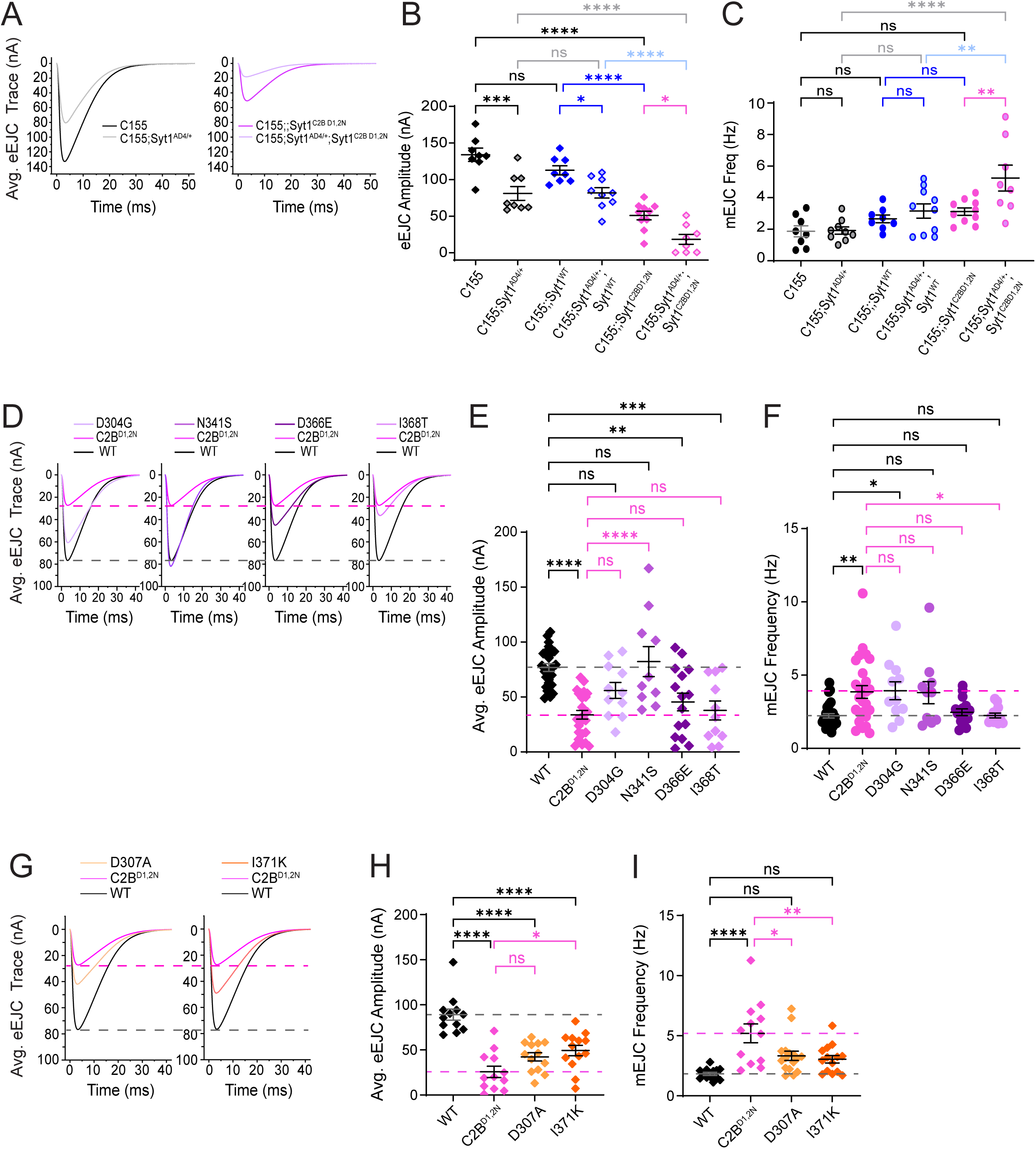
Disease-causing mutations in the SYT1 and SYT2 C2B Ca^2+^ binding loops reduce evoked release in a dominant manner. (A) Average eEJC traces for the indicated genotypes: *C155*, *C155;Syt1^AD4/+^*, *C155;;Syt1^C2BD1,2N^*, and *C155;Syt1^AD4/+^;Syt1^C2BD1,2N^*. (B) Quantification of mean eEJC amplitudes for the indicated genotypes. Note the decrease in release following *Syt1^C2BD1,2N^* overexpression is dosage-sensitive to endogenous SYT1 levels, with a ∼2.8-fold greater decrease in the *Syt1* null heterozygote than in animals with two wildtype *Syt1* alleles. (C) Quantification of mean spontaneous release rate for the indicated genotypes. Note *Syt1^C2BD1,2N^* overexpression only reduces SYT1’s clamping function in the *Syt1* null heterozygote background. (D) Representative eEJC responses for the indicated controls and SYT1 disease alleles: *C155;Syt1^AD4/+^;Syt1^WT^*(WT), *C155;Syt1^AD4/+^;Syt1^C2BD1,2N^ (C2B^D1,2N^), C155;Syt1^AD4/+^;Syt1^D304G^* (D304G), *C155;Syt1^AD4/+^;Syt1^N341S^* (N341S), *C155;Syt1^AD4/+^;Syt1^D366E^* (D366E) and *C155;Syt1^AD4/+^;Syt1^I368T^* (I368T). For comparison, the average WT trace is shown in black and its mean EJC amplitude represented by the dashed grey line. The C2B^D1,2N^ average trace is shown in magenta with its mean EJC amplitude represented by the dashed magenta line. (E) Quantification of mean eEJC amplitudes for the indicated genotypes. The WT mean eEJC amplitude is shown with the dashed grey line and the C2B^D1,2N^ mean eEJC amplitude with the dashed magenta line. Note the decreased evoked response following overexpression of the C2B Ca^2+^ binding pocket disease alleles compared with the non-pocket N341S allele. (F) Quantification of mean spontaneous release rate for the indicated genotypes. The dashed grey line shows the WT mean mEJC frequency and the dashed magenta line shows C2B^D1,2N^ mean mEJC frequency. (G) Representative eEJC responses for the indicated controls and SYT2 disease alleles (*C155;Syt1^AD4/+^;Syt1^WT^* (WT), *C155;Syt1^AD4/+^;Syt1^C2BD1,2N^ (C2B^D1,2N^), C155;Syt1^AD4/+^;Syt1^D307A^* (D307A) and *C155;Syt1^AD4/+^;Syt1^I371K^* (I371K). For comparison, the WT trace is shown in black with its mean EJC amplitude represented by the dashed grey line, while the C2B^D1,2N^ trace is shown in magenta with its mean EJC amplitude represented by a dashed magenta line. (H) Quantification of mean eEJC amplitudes for the indicated genotypes. The WT mean eEJC amplitude is shown with the dashed grey line and the C2B^D1,2N^ mean eEJC amplitude with the dashed magenta line. Note the decrease in evoked release following overexpression of the SYT2 C2B Ca^2+^ binding pocket disease alleles. (I) Quantification of mean spontaneous release rate for the indicated genotypes. The dashed grey line shows WT mean mEJC frequency and the dashed magenta line shows C2B^D1,2N^ mean mEJC frequency. Note that none of the disease alleles inhibit the clamping function of SYT2. Data are shown as mean ± SEM. Statistical significance: * *p* < 0.05, ** *p* < 0.005, *** *p* < 0.001, **** *p* < 0.0001, ns = not significant. Raw values, statistical tests and sample number are provided in the Source Data file.

We next examined the effects on synaptic transmission of disease-causing alleles in the C2B domain of SYT1 (D304G, N341S, D366E, I368T) and SYT2 (D307A, I371K) by mutating the corresponding residue within *Drosophila* SYT1. D304 and D366 correspond to the D1 and D4 Ca^2+^ binding residues within the SYT1 C2B domain, while I368 lies at the tip of one of the hydrophobic loops that penetrates the membrane upon Ca^2+^ binding (Figure 1C). N341 is not located in the C2B Ca^2+^ binding pocket but resides near the primary SYT1-SNARE binding interface on one side of the C2B domain. For SYT2 mutations, D307 corresponds to the D2 residue, while I371 sits in the base of the C2B Ca^2+^ binding pocket (Figure 1D). We generated UAS MYC-tagged transgenic lines for each of these mutations and wildtype SYT1 using site-specific transformation via the ΦC31 integrase system to eliminate genomic positional effects on transgene expression. Western analysis confirmed the mutant proteins were expressed at similar levels to the wildtype SYT1 transgenic protein except for the D307A allele that displayed a mild decrease (Supplemental Figure 1C, D, H, I). Immunostaining for SYT1 at NMJs demonstrated the mutant proteins undergo normal trafficking and accumulation within presynaptic terminals except for modestly reduced levels of D307A (Supplemental Figure 1E, G, J, L). Expression of most alleles did not alter synaptic growth at NMJs, with D366E and I368T causing a mild reduction in bouton number at 3^rd^ instar NMJs (Supplemental Figure 1F, K).

SYT1 and SYT2 transgenes that contained a mutation within the C2B Ca^2+^ binding pocket decreased evoked release when expressed in *Syt1^AD4/+^* heterozygotes (Figure 3D, E, G, H), with I368T representing the most severe allele and reducing SV fusion by >50%. Patients with the I368T mutation are also the most severe phenotypically, failing to develop language and displaying dystonia and self-injury^33,34^. In contrast to the disruption in evoked release, the majority of disease alleles did not alter spontaneous release frequency (Figure 3F, I). The N341S allele, which localizes to a distinct region of the SYT1 C2B domain, did not follow the pattern of mutations within the Ca^2+^ binding pocket. Expression of N341S did not reduce release when overexpressed, suggesting phenotypes of patients carrying this allele may result from a different mechanism (see below).

To test if disease alleles in the C2B Ca^2+^ binding pocket retain any ability to support synaptic transmission on their own, we expressed the mutants in the *Syt1* null background with *elav*^C155^ GAL4 and assayed evoked and spontaneous release in 3^rd^ instar larvae lacking any endogenous wildtype SYT1. Compared to rescue with wildtype SYT1, the D304G, D366E, I368T and D307A alleles failed to rescue synaptic release defects and caused a more severe impairment in evoked SV release than observed in the *Syt1* null background alone (Supplemental Figure 2A, B). For example, while complete loss of SYT1 decreased evoked SV fusion by 85% compared to controls, expression of the I368T allele reduced release by >96%. Most alleles also failed to rescue the enhanced spontaneous release rate observed in *Syt1* nulls (Supplemental Figure 2C). In summary, these data indicate alleles that disrupt the C2B Ca^2+^ binding pocket of both SYT1 and SYT2 dominantly disrupt evoked synaptic transmission when overexpressed in the presence of endogenous SYT1. In the absence of endogenous SYT1, they lack any ability to support evoked release on their own.

### Mutations in the C2B Ca^2+^ binding pocket are a privileged site for dominant-negative effects on synaptic transmission

Although Ca^2+^ binding to the C2B domain is a key trigger for activating SV fusion, other regions of the SYT1 protein play critical roles in regulating neurotransmitter release. To determine if mutations that alter other aspects of SYT1 function can act in a dominant-negative fashion, we generated transgenic lines expressing point mutations that disrupt other key interactions. Beyond Ca^2+^-dependent lipid binding, SYT1 interactions with the SNARE complex play a critical role in its ability to activate SV fusion^7,14,47–51^. The most essential site for this interaction is known as the primary SNARE binding interface that localizes to one side of the C2B domain and that can be disrupted by the SYT1 S280L mutation^7,51^. A secondary SYT1-SNARE interaction site within the C2B domain known as the tripartite interface includes CPX and can be disrupted with the L388Q, L395Q SYT1 allele^9,52^. In addition to SNARE interactions, SYT1 contain a polybasic stretch of amino acids on the opposite surface of the C2B domain that mediates Ca^2+^-independent lipid interactions to dock the protein on the PM^53–55^. The K327E mutation disrupts this interaction^7^. Finally, C2A-C2B intramolecular interactions help maintain the orientation of the membrane penetrating loops in the Ca^2+^-binding pockets and this intramolecular interaction can be disrupted with the R200H mutation^7^. UAS MYC-tagged transgenic proteins for each of these mutations were generated using the same genomic insertion site as above and overexpressed with *elav*^C155^ GAL4. Western analysis confirmed each protein was expressed at similar levels to the wildtype SYT1 transgenic version (Figure 4A, B). Unlike the strong reduction in evoked release caused by SYT1^C2BD1,2N^ and the C2B disease alleles, overexpression of SYT1^S280L^, SYT1^L388Q,L395Q^, SYT1^K327E^ or SYT1^R200H^ in *Syt1^AD4/+^* heterozygotes did not reduce evoked release or increase spontaneous fusion (Figure 4C-E). Together, these data indicate mutations disrupting the C2B Ca^2+^-binding pocket are likely to be the primary (if not only) region in which robust dominant-negative effects on synaptic transmission can be observed.

**Figure 4.**
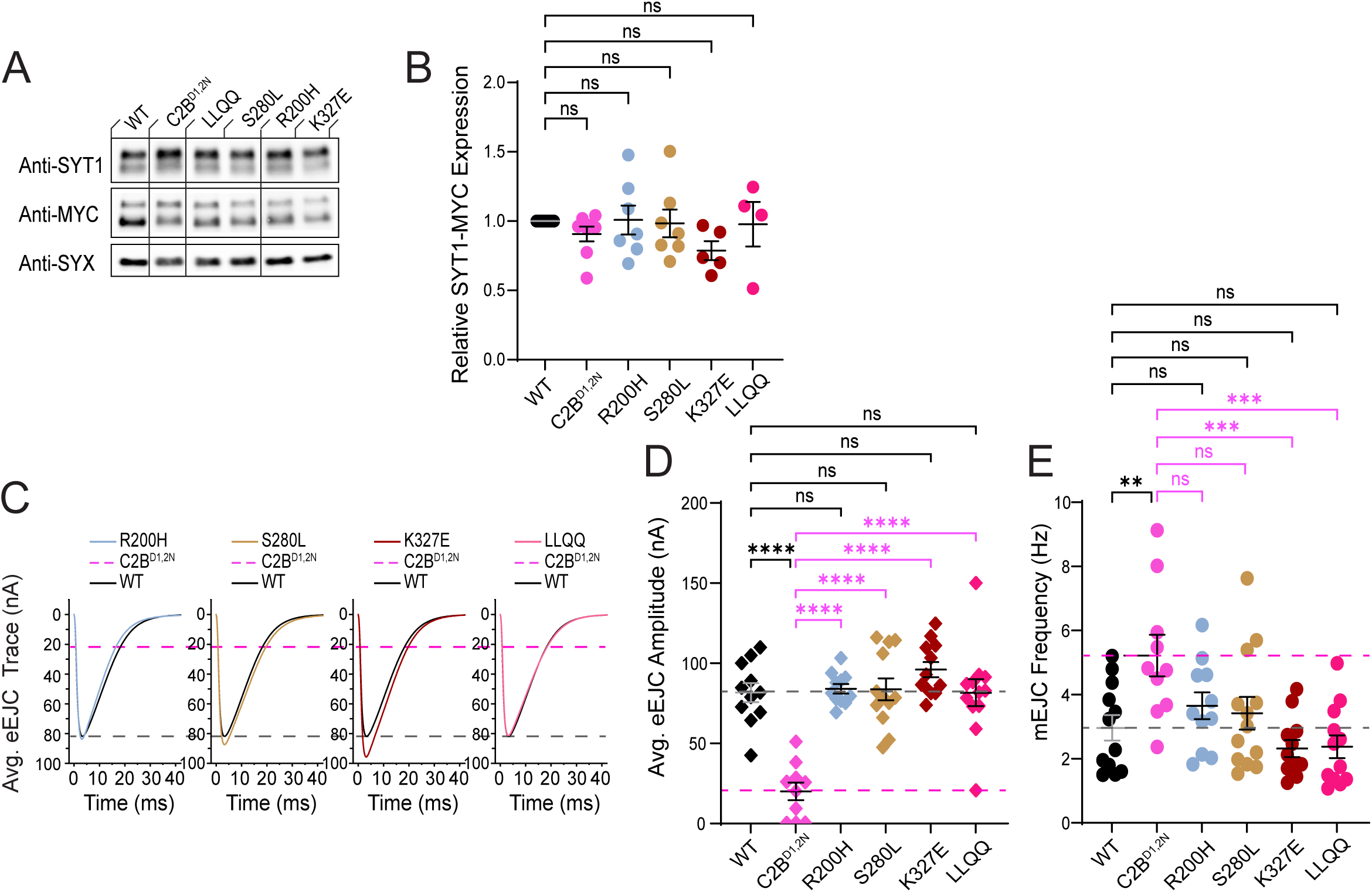
Mutations in the C2B Ca^2+^ binding pocket are a privileged site for dominant-negative effects on synaptic transmission. (A) Representative western for SYT1 (anti-SYT1) and SYT1-MYC expression (anti-MYC) from adult brain extracts with anti-SYX as a loading control for the indicated genotypes: *C155;Syt^AD4/+^;Syt^WT^*(WT), *C155;Syt^AD4/+^;Syt^C2BD1,2N^* (C2B^D1,2N^), *C155;Syt^AD4/+^;Syt^L388Q,L395Q^* (LLQQ), *C155;Syt^AD4/+^; Syt^S280L^* (S280L), *C155;Syt^AD4/+^;Syt^R200H^*(R200H), and *C155;Syt^AD4/+^;Syt^K327E^* (K327E). (B) Quantification of SYT1-MYC protein expression normalized to control (WT) for the indicated genotypes. (C) Average eEJC traces for the indicated genotypes. For comparison, the WT mean eEJC amplitude is shown with a dashed grey line and the C2B^D1,2N^ mean eEJC amplitude with a dashed magenta line. (D) Quantification of mean eEJC amplitudes for the indicated genotypes. The WT mean eEJC amplitude is shown with the dash grey line and the C2B^D1,2N^ mean eEJC amplitude with the dashed magenta line. Note that none of these mutations in SYT1 decrease the evoked response when overexpressed. (E) Quantification of mean spontaneous release rate for the indicated genotypes. The dashed grey line represents WT mean mEJC frequency and the dashed magenta line shows C2B^D1,2N^ mean mEJC frequency. Data are shown as mean ± SEM. Statistical significance: ** *p* < 0.005, *** *p* < 0.001, **** *p* < 0.0001, ns = not significant. Raw values, statistical tests and sample number are provided in the Source Data file.

### Mutations in the C2B Ca^2+^ binding pocket cause dominant-negative effects regardless of changes to electrostatic charge in the region

Ca^2+^ binding to SYT’s C2 domains alters the electrostatic potential near the membrane binding loops by neutralizing negative charges from the five aspartic acid residues and allowing nearby hydrophobic and basic residues to insert into anionic lipids of the PM. One hypothesis for the dominant effects of C2B Ca^2+^ binding alleles is that they reduce negative charge in this region to mimic Ca^2+^ binding, allowing the C2B domain loops to insert into the PM in the absence of Ca^2+^ to disrupt evoked release. To test this model, we generated transgenic lines that alter or preserve the negative charge of aspartic acid residues in C2B, similar to prior studies testing a role for an electrostatic switch in the C2A and C2B Ca^2+^ binding pockets^56,57^. For this analysis, we targeted the C2B D4 aspartic acid given one disease allele alters this residue (D366). We mutated C2B D4 to preserve the negative charge (D4E), neutralize the negative charge (D4N) or reverse the charge to a positive residue (D4R). If disease alleles act in part by causing the C2B domain to insert into the lipid bilayer in a Ca^2+^-independent manner, we hypothesized C2B^D4N^ where the charge is neutralized, and C2B^D4R^ where a positive charge is added to enhance anionic lipid binding, would display more severe disruptions to neurotransmitter release than C2B^D4E^. Each mutant allele was expressed at similar levels as the wildtype SYT1 protein (Figure 5A, B), indicating these residue substitutions do not affect SYT1 stability. The decrease in evoked release caused by expression of the C2B^D4^ mutants in *Syt1^AD4/+^* heterozygotes were not significantly different from each other (Figure 5C, D). Similar to other disease alleles, spontaneous release frequency was not affected, indicating a selective defect in SYT1’s Ca^2+^-dependent triggering of evoked release compared to the Ca^2+^-independent clamping function that requires SNARE binding^7^. We conclude that the dominant effects of the C2B Ca^2+^ binding pocket disease alleles are unlikely to be secondary to electrostatic potential changes that trigger Ca^2+^-independent membrane insertion by the C2B loops.

**Figure 5.**
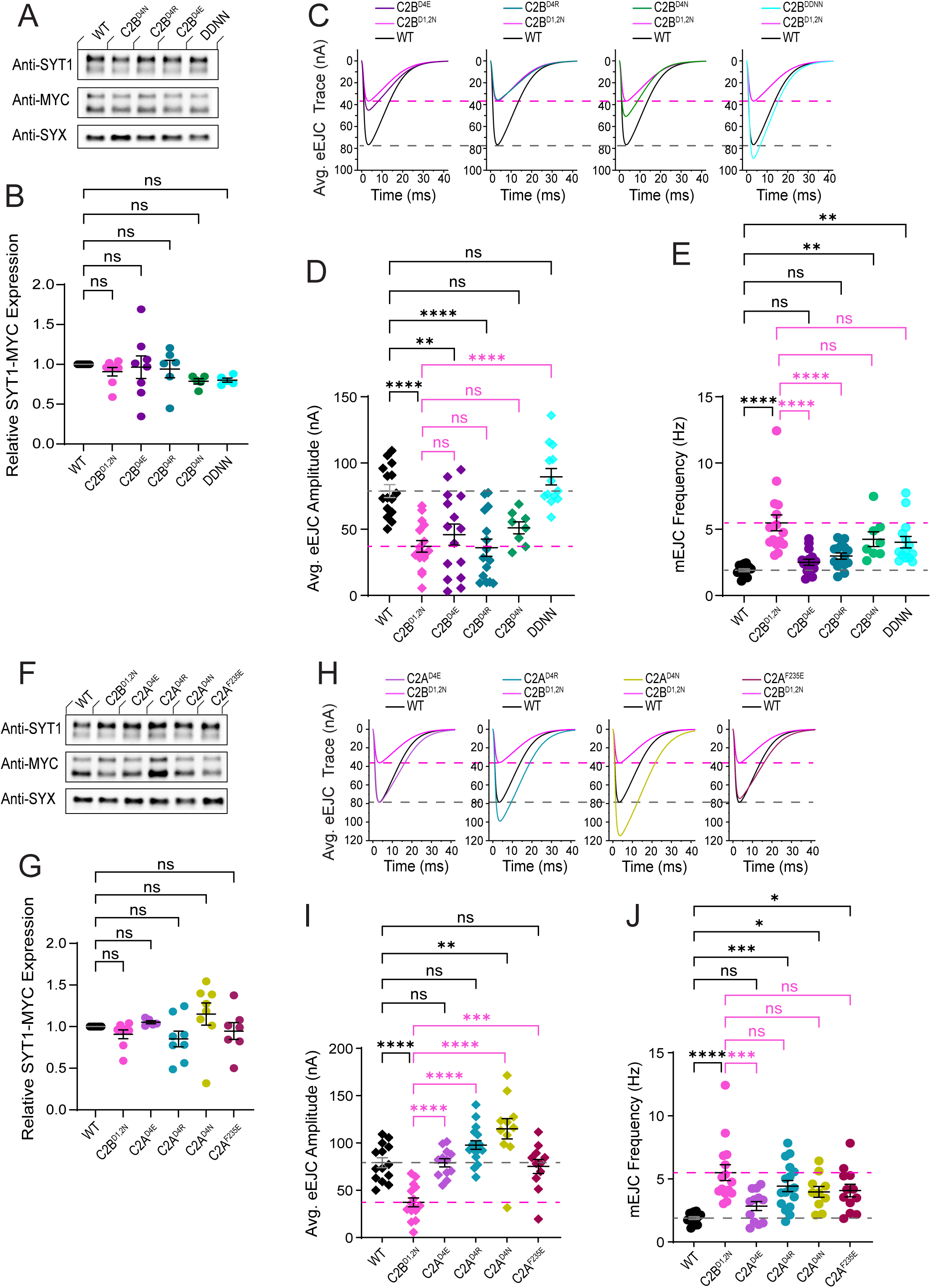
Overexpression effects of charge-altering mutations in the SYT1 C2A and C2B Ca^2+^ binding pockets. (A) Representative western for SYT1 (anti-SYT1) and SYT1-MYC expression (anti-MYC) from adult brain extracts with anti-SYX as a loading control for the indicated genotypes: *C155;Syt1^AD4/+^;Syt1^WT^* (WT), *C155;Syt1^AD4/+^;Syt1^C2BD1,2N^* (C2B^D1,2N^), *C155;Syt1^AD4/+^;Syt1^C2B-D4E^*(C2B^D4E^), *C155;Syt1^AD4/+^;Syt1^C2B-D4R^* (C2B^D4R^), *C155;Syt1^AD4/+^;Syt1^C2B-D4N^* (C2B^D4N^), and *C155;Syt1^AD4/+^;Syt1^C2A-D4N,C2B-D4N^* (DDNN). (B) Quantification of SYT1-MYC protein expression normalized to control (WT) for the indicated genotypes. (C) Average eEJC traces for the indicated genotypes described. For comparison, the WT average trace is shown in black and its mean EJC amplitude with the dashed grey line. The C2B^D1,2N^ average trace is shown in magenta and its mean EJC amplitude with the dashed magenta line. (D) Quantification of mean eEJC amplitudes for the indicated genotypes. The WT mean eEJC amplitude is shown with the dashed grey line and the C2B^D1,2N^ mean eEJC amplitude with the dashed magenta line. Note that the C2B Ca^2+^ pocket mutations decrease evoked release when overexpressed, while double mutation of both the C2A and C2B Ca^2+^ binding domains block the dominant effect. (E) Quantification of mean spontaneous release rate for the indicated genotypes. The dashed grey line represents WT mean mEJC frequency and the dashed magenta line shows C2B^D1,2N^ mean mEJC frequency. (F) Representative western for SYT1 (anti-SYT1) and SYT1-MYC expression (anti-MYC) from adult brain extracts with anti-SYX as a loading control for the indicated genotypes: *C155;Syt1^AD4/+^;Syt1^WT^* (WT), *C155;Syt1^AD4/+^;Syt1^C2BD1,2N^* (C2B^D1,2N^), *C155;Syt1^AD4/+^;Syt1^C2A-D4E^*(C2A^D4E^), *C155;Syt1^AD4/+^;Syt1^C2A-D4R^* (C2A^D4R^), *C155;Syt1^AD4/+^;Syt1^C2A-D4N^* (C2A^D4N^), and *C155;Syt1^AD4/+^;Syt1^F235E^*(C2A^F235E^). (G) Quantification of SYT1-MYC protein expression normalized to control (WT) for the indicated genotypes. (H) Average eEJC traces for the indicated genotypes. For comparison, the WT average trace is shown in black and its mean EJC amplitude is represented by the dashed grey line. The C2B^D1,2N^ average trace is shown in magenta with its mean EJC amplitude represented by the dashed magenta line. (I) Quantification of mean eEJC amplitudes for the indicated genotypes. The WT mean eEJC amplitude is shown with the dashed grey line and the C2B^D1,2N^ mean eEJC amplitude with the dashed magenta line. Note that overexpression of the C2A Ca^2+^ pocket alleles do not decrease evoked release, with C2A^D4R^ and C2A^D4N^ actually increasing release. (J) Quantification of mean spontaneous release rate for the indicated genotypes. The WT mean mEJC frequency is shown with the dashed grey line and the C2B^D1,2N^ mean mEJC frequency is shown with the dashed magenta line. Data are shown as mean ± SEM. Statistical significance: * *p* < 0.05, ** *p* < 0.005, *** *p* < 0.001, **** *p* < 0.0001, ns = not significant. Raw values, statistical tests and sample number are provided in the Source Data file.

### Mutations in the C2A Ca^2+^ binding pocket do not cause dominant-negative effects on their own but prevent toxicity of the C2B Ca^2+^ binding pocket mutants

Like C2B, the C2A domains of SYT1 and SYT2 contain five negatively charged aspartic acid residues embedded within their two membrane-penetrating loops. To date, no disease-causing mutations in human SYT1 or SYT2 localize to the C2A Ca^2+^ binding pocket. To determine if mutations disrupting Ca^2+^ binding to C2A could dominantly disrupt neurotransmitter release, we generated transgenic lines expressing SYT1 with the C2A D4 residue mutated (C2A^D4N^, C2A^D4R^, and C2A^D4E^) as described above for the C2B D4 alleles. We also generated a mutation (C2A^F235E^) previously shown to block Ca^2+^-dependent membrane penetration by the C2A domain^13^. This allele changes a hydrophobic phenylalanine residue in a C2A membrane penetration loop to a negatively charged glutamic acid that repels anionic phospholipids. All four C2A alleles were expressed at the same level as wildtype SYT1 (Figure 5F, G). Unlike the C2B^D4^ alleles that decrease evoked release (Figure 5D), the same reside changes in C2A did not reduce synaptic transmission when overexpressed in *Syt1^AD4/+^* heterozygotes (Figure 5H, I). Instead, C2A^D4N^ and C2A^D4R^, which reduce the negative charge in the C2A Ca^2+^ binding pocket and are predicted to enhance C2A membrane penetration, resulted in increased evoked and spontaneous SV release (Figure 5H-J). This phenotype is consistent with a prior study from the lab demonstrating *Syt1* transgenes containing a C2A^D3N,D4N^ mutation enhance release when expressed in the *Syt1* null background^20^.

Although mutations in the C2A Ca^2+^ binding pocket did not decrease release when overexpressed, it is possible C2A Ca^2+^ binding and membrane penetration is required for the toxicity of C2B mutants, as both C2 domains penetrate the lipid bilayer during SV fusion. To test this model, we generated a transgenic line disrupting the function of both C2A and C2B Ca^2+^ binding pockets (C2A^D4N^, C2B^D4N^). The dual C2 mutant (DDNN) protein was expressed at similar levels as wildtype SYT1 (Figure 5A, B). Unlike the reduction in evoked release caused by the C2B^D4N^ mutation alone, overexpression of C2A^D4N^, C2B^D4N^ in *Syt1^AD4/+^* heterozygotes did not act in a dominant manner to decrease evoked release (Figure 5C, D). We conclude that mutations that disrupt Ca^2+^ binding (C2A^D4N^, C2A^D4R^, and C2A^D4E^) or membrane penetration (C2A^F235E^) by the C2A domain do not act in a dominant-negative manner, in contrast to similar mutations in the C2B domain. However, the normally dominant-negative effects of C2B Ca^2+^ binding pocket mutants are prevented by simultaneous loss of C2A Ca^2+^-dependent lipid binding. These data indicate C2A membrane insertion is likely required for proper positioning at fusion sites for the mutant C2B domain to poison the fusion machinery, similar to the requirement for SNARE and C2B Ca^2+^-independent membrane binding^7^.

### Molecular dynamics simulation of C2B Ca^2+^ binding pocket alleles reveals defects in membrane insertion

To examine how mutations in the C2B Ca^2+^ binding pocket disrupt neurotransmitter release by altering Ca^2+^-dependent interactions with the PM, molecular dynamics (MD) simulation was performed to compare wildtype SYT1 with disease alleles (D304G, D366E, I368T for SYT1 and D307A, I371K for SYT2). Initial runs of 4.8 µs were performed to follow attachment of the Ca^2+^-free C2B domain to a POPC:POPS:PIP_2_ bilayer mimicking the PM (Figure 6A, B). Within 0.5 µs, each variant attached to the PM and remained attached until the end of the trajectory. As previously observed^58^, the C2B domain attached to PIP_2_ monomers in the PM via three anchors, including the polybasic stretch (PB), basic residues within the Ca^2+^-binding loops (CBL), and a RR motif at the base of the C2B domain (Figure 6A). These anchors produced a roughly parallel orientation of the C2B domain against the PM with extensive C2B-PM surface interactions. The D307A, D366E, and I368T alleles attached to the PM in a similar fashion to wildtype, while D304G and I371K were tilted with only the PB and CBL anchors attaching to the PM. At the end of the MD trajectory, the five Ca^2+^ binding aspartate residues in wildtype SYT1 clustered together and were exposed to solution in a pattern predicted to facilitate Ca^2+^ chelation. This arrangement was altered in the SYT disease alleles (Figure 6B). Mutation of the D2 residue (D307A) caused the D1 residue in loop 1 to separate from the cluster, while mutation of D1 (D304G) caused the D3 and D4 residues in loop 3 to be shielded by loop 1 due to the tilted orientation of the protein. I371K had a similarly tilted orientation, with the D3 and D4 residues in loop 3 shielded by loop 1 and lipids. Although the acidic residues clustered together in the D4 (D366E) allele, the D3 residue in loop 3 was shielded by the aspartate to glutamate substitution. Finally, the I368T mutant resulted in separation of the two loops, distancing the D1 residue in loop 1 from the remainder of the cluster.

**Figure 6.**
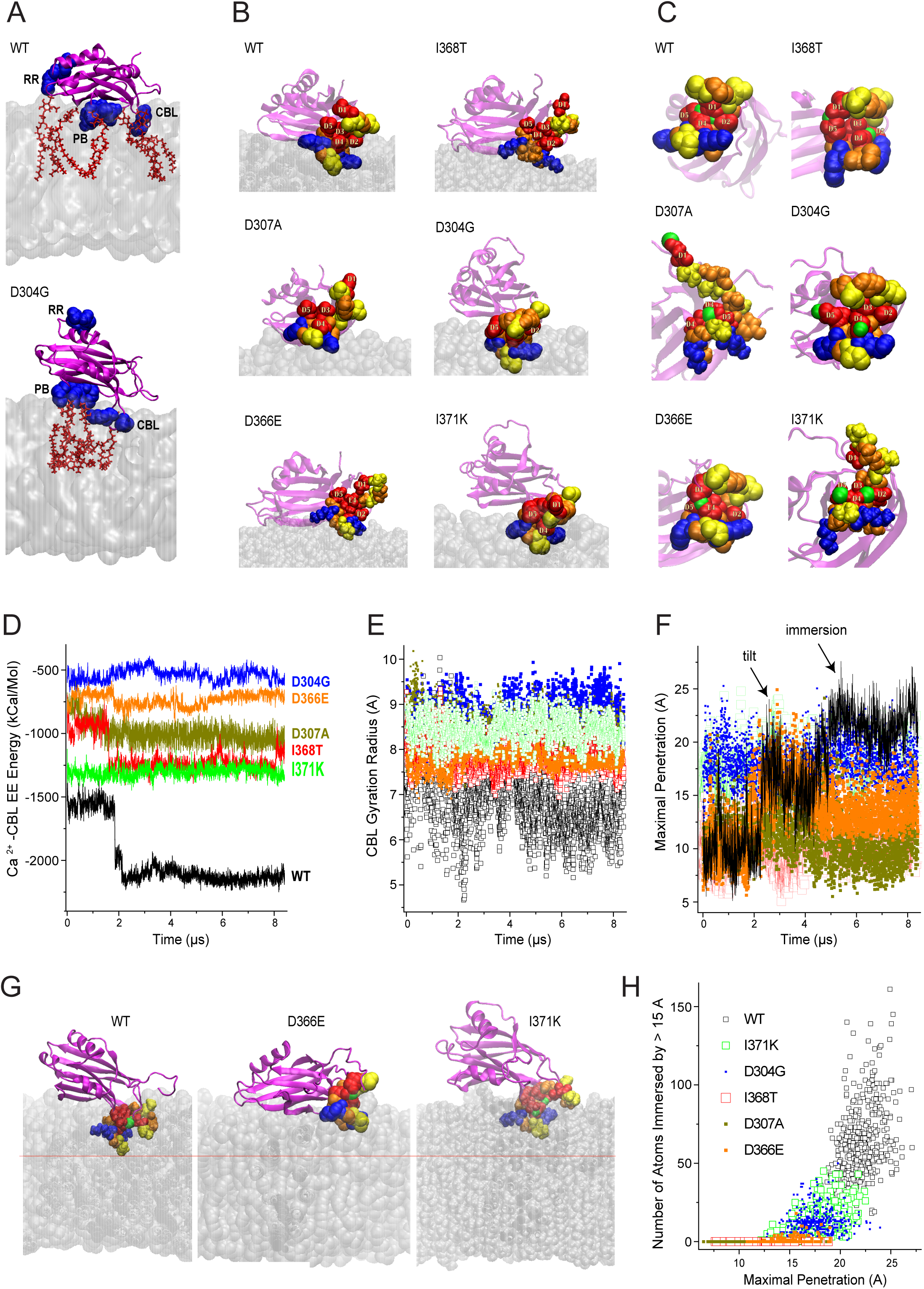
MD simulation of Ca^2+^ binding and membrane penetration by control SYT1 and disease alleles. (A) Attachment of the WT SYT1 C2B domain to the lipid bilayer (silver, quick surface representation) at the end of the 4.8 µs MD trajectory. SYT1 is attached to PIP_2_ monomers (red, bond representation) via three anchors (blue, VdW representation): two basic residues of the Ca^2+^-binding loop 3 (CBL), the polybasic stretch (PB), and the RR motif. In contrast, the D304G allele is tilted and attached to the PM through only two anchors (CBL and PB). (B) Arrangement of the Ca^2+^ binding residues at the end of the 4.8 µs MD trajectories. Ca^2+^ - binding loops 1 and 3 are shown in WdV representation: red - acidic; blue – basic; yellow – hydrophobic; orange – polar. Note the tight symmetric arrangement of the D1-D5 residues in WT SYT1 compared to the disrupted pattern in all the disease alleles. (C) Ca^2+^ coordination at the end of the 8.4 µs MD trajectories. Ca^2+^ ions (green) are shown in VdW representation. Note that two Ca^2+^ ions are cooperatively chelated by all the aspartic residues in WT SYT1. This coordination is disrupted in all the disease alleles. (D) Electrostatic energy of the interactions of Ca^2+^ ions with the Ca^2+^-binding loops over the length of the MD trajectories. Note a sharp drop in the energy in WT SYT1 around the 2 µs time-point, which does not occur in the disease alleles. (E) Gyration radius computed for the residues forming the Ca^2+^-binding loops. The color coding for WT (black) and the mutants is shown in panel D. Note the drop in gyration radius in the second part of the trajectory for WT SYT1, indicating that the loops tightened. (F) Maximal atomic penetration into lipids during the MD trajectories. Note two conformational transitions for the WT protein (arrows). The second transition (immersion) did not occur in any of the disease alleles. (G) Structures of WT and two disease alleles (D366E and I371K) at the end of their respective trajectories. The disease alleles do not immerse into the PM to the same level compared to WT SYT1 (marked with the orange line). The PM (silver) is shown in the VdW representation. (H) Penetration level measured in the space of two variables: the number of atoms of the protein immersed by over 15 Å into the bilayer versus the maximal atomic penetration. Data was collected from the last microsecond (7.4 to 8.4 µs) of the respective trajectories. Each data point represents a single trajectory point. Note the cluster of the WT datapoints (black) shifted towards the larger penetration values, which does not overlap with the clusters observed for disease alleles.

The observed changes in the pattern of C2B Ca^2+^-binding residues suggest Ca^2+^ chelation and membrane penetration may be altered by the SYT disease alleles. To test this model, two Ca^2+^ ions were added to each system by K^+^ replacements as described^59^. For each C2B-Ca^2+^ complex attached to the PM, a 8.4 µs MD run was performed and Ca^2+^ coordination and PM penetration were analyzed (Figure 6C-E). For all variants except D366E, two Ca^2+^ ions were chelated within the Ca^2+^ binding pocket for the entire trajectory (Figure 6C). D366E retained only a single Ca^2+^ ion that was chelated simultaneously by multiple acidic residues from both loops. The second Ca^2+^ ion was rapidly lost from the pocket and did not attach to any residues in the Ca^2+^-binding loops over the length of the trajectory. Although four out of five C2B alleles retained two Ca^2+^ ions in the Ca^2+^-binding pocket, Ca^2+^ coordination of the C2B domain was distinct for all the disease alleles compared to control (Figure 6C). For wildtype SYT1, two Ca^2+^ ions were cooperatively chelated by the five aspartate residues in a symmetric fashion, with each of the Ca^2+^ ions forming four coordination bonds (Figure 6C). D1, D3, and D4 formed two coordination bonds (one with each of the two Ca^2+^ ions), while D2 and D5 formed a single coordination bond. This Ca^2+^ binding configuration was formed within the initial 2 µs and remained intact until the end of the 8.4 µs trajectory (Figure 6D). After the Ca^2+^ ions were cooperatively chelated, the loops became tighter, as illustrated by their diminished gyration radius along the trajectory (Figure 6E). None of the disease alleles displayed similar Ca^2+^ coordination, with overall electrostatic interactions of Ca^2+^ ions with the Ca^2+^-binding loops drastically weakened compared to control (Figure 6D). The disease alleles failed to show any decrease in the gyration radius of the loops (Figure 6E), with the Ca^2+^ coordination and organization of the loops differing for each variant. For wildtype SYT1, both Ca^2+^ ions were enclosed in the pocket, with hydrophobic residues of loop 1 (V305 and L308) and loop 3 (I368) approaching each other and shielding the negatively charged acidic residues (Figure 6C). In the D366E variant, the overall organization of the loops was similar, with a single Ca^2+^ ion chelated by one aspartic residue of loop 1 and two residues (Asp and Glu) in loop 3, bringing the two loops together. However, the D366E substitution shielded the remaining two aspartic residues from chelating a second Ca^2+^ ion. The I368T variant had one of the two Ca^2+^ ions chelated solely by residues in loop 3, while the other Ca^2+^ ion was able to bridge loops 1 and 3. In addition, the D366E variant lacked one Ca^2+^ ion as noted above, while I368T lacked a hydrophobic residue in the Ca^2+^ binding tip. For the D304G, D307A and I371K alleles, the loops were not bridged and each Ca^2+^ ion was chelated by either a single loop or by a single aspartic acid (Figure 6C). Thus, all the variants had an altered pattern of Ca^2+^ binding, resulting in an abnormal structure for the Ca^2+^ binding pocket that caused its hydrophobic residues to be more dispersed.

We next analyzed the trajectories of the Ca^2+^-bound variants to evaluate the depth of PM penetration (Figure 6F-H). Two parameters were used to assay PM penetration: maximal penetration depth, measured as the distance of the most immersed atom to the surface of the bilayer (Figure 6F); and the number of atoms immersed by more than 15 Å under the bilayer surface (Figure 6H). At the end of the trajectory, wildtype SYT1 had the entire Ca^2+^-bound tip, including loops 1 and 3, immersed within the bilayer (Figure 6F). The protein underwent two conformational transitions along the trajectory, with the protein tilting after 2 µs to produce immersion of loop 1, and a subsequent immersion of loop 3 after 4 µs (Figure 6F). None of the disease alleles underwent these conformational transitions and showed little or no Ca^2+^-dependent PM penetration (Figure 6F). The only exception was the D366E allele, which showed a tilt after 2 µs followed by a subsequent reversal after 4 µs. At the end of the trajectory, D366E mostly displayed surface interactions with the lipid bilayer, with loop 3 being partially immersed and loop 1 remaining in solution (Figure 6G). Similar levels of reduced PM penetration were observed for the I368T and D307A variants. The D304G and I371K variants, which showed immersion of loop 3 with the PM in their Ca^2+^-free form, retained the same conformation with loop 3 being immersed and loop 1 showing only surface interactions (Figure 6F-H). In summary, MD simulations indicate the C2B Ca^2+^-binding domain is fine-tuned to allow acidic residues in both loops to cooperatively chelate two Ca^2+^ ions to promote a closed loop conformation. This enables hydrophobic loop residues to penetrate into the lipid bilayer, leading to immersion of the entire Ca^2+^-bound tip of the C2B domain into the PM. None of the disease alleles were capable of PM immersion to the same level as observed for wildtype SYT1, consistent with disrupted Ca^2+^ coordination and distorted structure of the Ca^2+^-bound loops caused by these mutations.

### Disease alleles outside of the C2B Ca^2+^ binding pocket act through haploinsufficient or gain-of-function mechanisms

We next examined how disease alleles in SYT1 outside of the C2B Ca^2+^ binding pocket alter neurotransmitter release, including the L159R, T196K, E209K and E219Q C2A and N341S C2B variants that cause less severe patient phenotypes (Figure 1A-C). UAS MYC-tagged transgenic lines for each of these mutations was generated as described above and compared to the effects of wildtype SYT1 or dominant-negative SYT1C2B^D1,2N^. Unlike the C2B Ca^2+^ pocket alleles, western analysis indicated the L159R and T196K mutations reduced SYT1 protein levels compared to the wildtype SYT1 transgenic protein or the E209K, E219Q and N341S alleles (Figure 7A, B, Supplemental Figure 3A). Given the L159R residue change leads to an unstable protein that was fully degraded and undetectable by western, we did not perform additional analysis of this allele. Immunostaining with anti-MYC demonstrated the remaining SYT1 alleles underwent normal trafficking and accumulation within synaptic terminals except for a ∼72% reduction in presynaptic T196K levels (Figure 7C, D), consistent with its reduced protein expression. Synaptic growth at larval NMJs was not affected following overexpression of these alleles (Figure 7E). Compared to SYT1C2B^D1,2N^, these alleles did not alter evoked (Figure 7F, G) or spontaneous (Figure 7H) release when overexpressed in *Syt1^AD4/+^*heterozygotes with *elav*^C155^ GAL4. We conclude these point mutants do not dominantly block SV fusion as observed for the C2B Ca^2+^ binding alleles.

**Figure 7.**
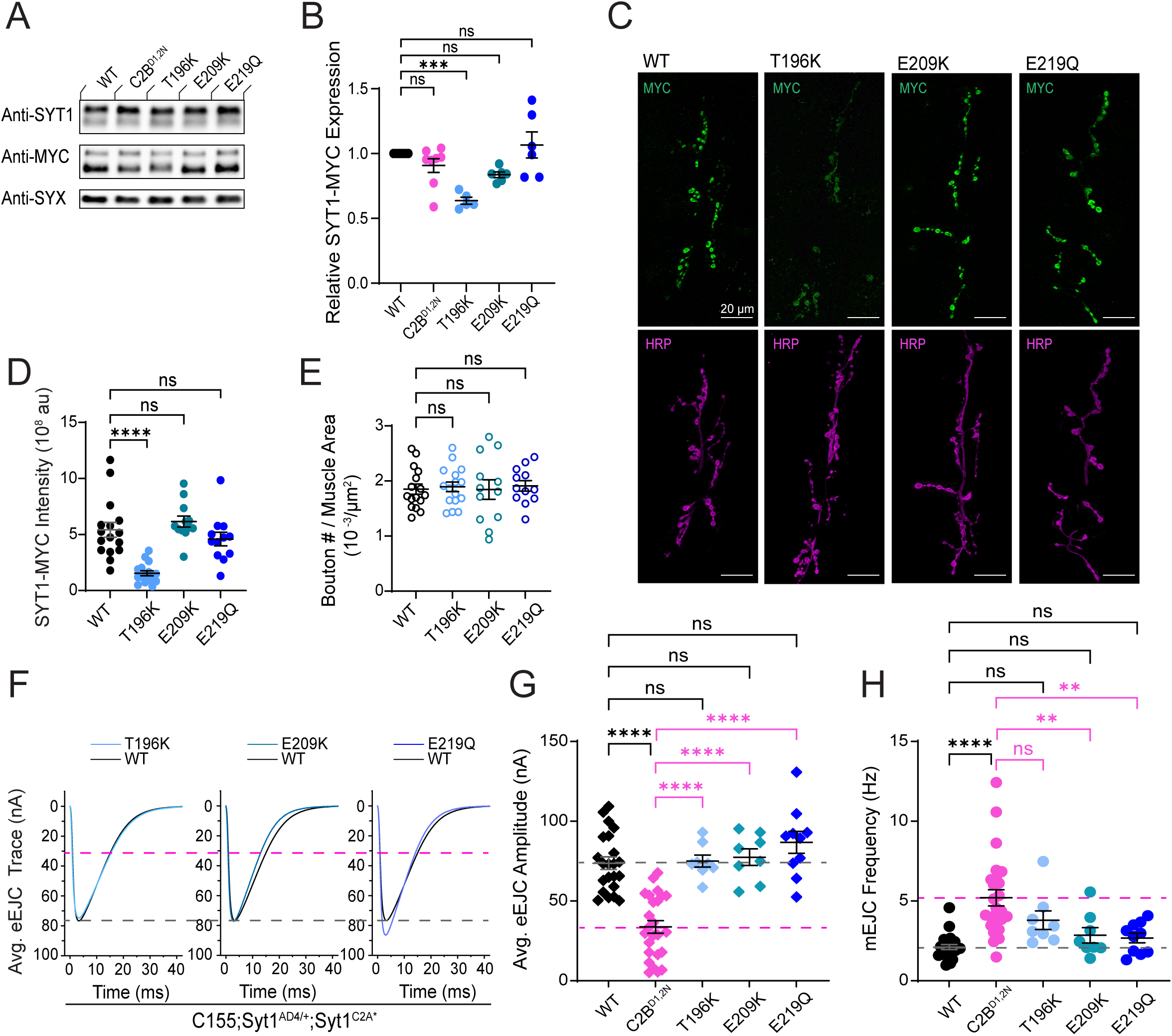
Disease-causing mutations in the SYT1 C2A domain do not reduce evoked release in a dominant manner. (A) Representative western for SYT1 (anti-SYT1) and SYT1-MYC expression (anti-MYC) from adult brain extracts with anti-SYX as a loading control for the indicated genotypes: *C155;Syt1^AD4/+^;Syt1^WT^*(WT), *C155;Syt1^AD4/+^;Syt1^C2BD1,2N^* (C2B^D1,2N^), *C155;Syt1^AD4/+^;Syt1^T196K^* (T196K)*, C155;Syt1^AD4/+^;Syt1^E209K^*(E209K), and *C155;Syt1^AD4/+^;Syt1^D219Q^* (D219Q). (B) Quantification of SYT1-MYC protein expression normalized to control (WT) for the indicated genotypes. Note the reduced protein levels of the T196K allele. (C) Immunostaining for SYT1 (anti-MYC, green) and HRP (magenta) at 3^rd^ instar muscle 6/7 NMJs of the indicated genotypes. (D) Quantification of total SYT1-MYC fluorescence within the HRP-positive NMJ area (au = arbitrary units). Note the dramatic reduction in synaptic levels of the T196K allele. (E) Quantification of varicosity number normalized to muscle surface area for the indicated genotypes. (F) Average eEJC traces for the indicated genotypes. The WT mean eEJC amplitude is shown with a dashed grey line and the C2B^D1,2N^ mean eEJC amplitude with a dashed magenta line. (G) Quantification of mean eEJC amplitudes for the indicated genotypes. The WT mean eEJC amplitude is shown with a dashed grey line and the C2B^D1,2N^ mean eEJC amplitude with a dashed magenta line. Note that none of these mutations in SYT1 decrease the evoked response when overexpressed. (H) Quantification of mean spontaneous release rate for the indicated genotypes. WT mean mEJC frequency is shown with the dashed grey line and C2B^D1,2N^ mean mEJC frequency with the dashed magenta line. Overexpression of these alleles does not alter SYT1’s clamping function. Data are shown as mean ± SEM. Statistical significance: ** *p* < 0.005, *** *p* < 0.001, **** *p* < 0.0001, ns = not significant. Raw values, statistical tests and sample number are provided in the Source Data file.

To determine if this group of mutations alters the endogenous function of SYT1, the alleles were expressed in the *Syt1* null background with *elav*^C155^ GAL4. Western analysis confirmed the mutant proteins were expressed at similar levels to the wildtype SYT1 transgenic protein except for the T196K allele (Figure 8A, B). Immunostaining for SYT1-MYC at NMJs demonstrated the proteins underwent normal trafficking and accumulation within presynaptic terminals except for T196K, which showed a 83% reduction in presynaptic levels in the null background (Figure 8C, D). Expression of these alleles did not alter synaptic growth at larval NMJs (Figure 8E). To determine their functional properties, evoked and spontaneous release was quantified in 3^rd^ instar rescued larvae expressing the mutant allele and lacking any endogenous SYT1. Compared to rescue with wildtype SYT1, only the E209K allele showed a normal ability to support synaptic transmission (Figure 8F-H). Given E209K can drive normal levels of neurotransmitter release on its own and does not reduce synaptic transmission in a dominant fashion, it is possible this allele is non-pathogenic or requires additional synergistic mutations specific to the patient’s genetic background. The T196K allele completely failed to rescue the reduced evoked release observed in the *Syt1* null background, consistent with its 83% reduction in presynaptic protein abundance. We conclude that the decreased stability of the L159R and T196K SYT1 proteins cause a haplo-insufficient phenotype that reduces synaptic transmission as observed in *Syt1* mutant heterozygotes (Figure 2). In contrast to the inability of T196K to rescue synaptic transmission in the *Syt1* null background, the E219Q and N341S alleles resulted in a 3.1 and 5-fold increase in evoked release compared to rescue with wildtype SYT1 (Figure 8F, G). In addition to the effect on evoked release, the N341S allele increased the frequency of spontaneous release by 6.5-fold (Figure 8H). These data indicate the E219Q allele enhances the ability of SYT1 to active evoked fusion, while N341S increases both evoked and spontaneous SV release. We conclude these two alleles act through a gain-of-function mechanism to increase SYT1’s ability to activate SV fusion that results in excessive neurotransmitter release.

**Figure 8.**
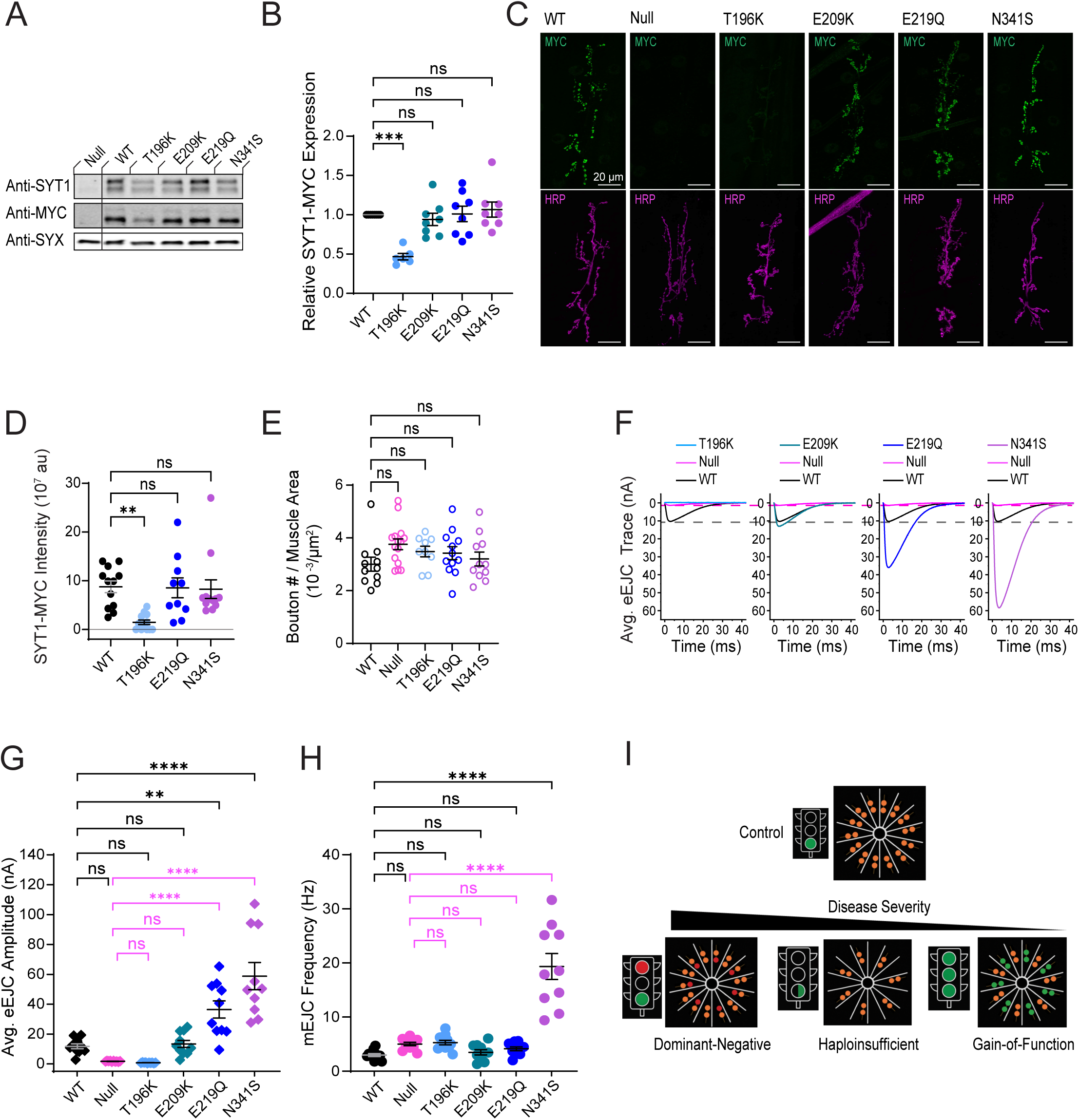
SYT1 disease alleles outside of the C2B Ca^2+^ binding pocket can reduce SYT1 expression or enhance SYT1 function. (A) Representative western for SYT1 (anti-SYT1) and SYT1-MYC expression (anti-MYC) from adult brain extracts with anti-SYX as a loading control for the indicated genotypes: *C155;Syt1^AD4/N13^* (Null mutant), *C155;Syt1^AD4/N13^;Syt1^WT^* (WT), *C155;Syt1^AD4/N13^;Syt1^T196K^* (T196K), *C155;Syt1^AD4/N13^;Syt1^E209K^* (E209K), *C155;Syt1^AD4/N13^;Syt1^E219Q^* (E219Q), and *C155;Syt1^AD4/N13^;Syt1^N341S^* (N341S). (B) Quantification of SYT1-MYC protein expression normalized to rescue with WT SYT1 for the indicated genotypes. Note the reduced protein levels of the T196K allele. (C) Immunostaining for SYT1 (anti-MYC, green) and HRP (magenta) at 3^rd^ instar muscle 6/7 NMJs for the indicated genotypes. (D) Quantification of total SYT1-MYC fluorescence within the HRP-positive NMJ area (au = arbitrary units). Note the dramatic reduction in synaptic levels of the T196K allele. (E) Quantification of varicosity number normalized to muscle surface area for the indicated genotypes. (F) Average eEJC traces for the indicated genotypes. The mean EJC trace for rescue of the null mutant with WT SYT1 is shown in black and its mean EJC amplitude represented by the dashed grey line. The average mean EJC trace for the *Syt1* null mutant is shown in magenta with its mean EJC amplitude represented by the dashed magenta line. (G) Quantification of mean eEJC amplitudes for the indicated genotypes. Note the failure of the T196K allele to rescue and the enhanced rescue with the E219Q and N341S alleles. (H) Quantification of mean spontaneous release rate for the indicated genotypes. Note the dramatic enhancement in spontaneous release with the N341S rescue. Data are shown as mean ± SEM. Statistical significance: ** *p* < 0.005, *** *p* < 0.001, **** *p* < 0.0001, ns = not significant. Raw values, statistical tests and sample number are provided in the Source Data file. (I) Model for disease severity across the SYT1 alleles. SNARE complexes are shown as spokes on a wheel at the SV-PM interface with SYT1 (orange) bound to each SNARE complex. In the control case, Ca^2+^ binding by SYT1 acts as go signal to trigger SNARE-dependent fusion. For the most severe dominant-negative Ca^2+^ binding loop alleles, half of the SYT1 proteins will not be able to undergo Ca^2+^ -dependent PM insertion due to the mutation in C2B (red circles), resulting in poisoning of release sites. For haploinsufficient alleles, the 50% reduction in SYT1 protein levels results in a less efficient go signal and decreased SV release. In contrast, gain-of-function alleles (green) result in excessive SYT1 function and enhanced SV fusion.

## Discussion

SYT1 and SYT2 have multiple roles in regulating the SV cycle, including SV priming, Ca^2+^-triggering of fast synchronous release, suppression of asynchronous release, clamping spontaneous fusion, and promoting SV endocytosis^1,6^. While SYT2 expression and function is concentrated at peripheral NMJs, SYT1 is abundant in the CNS. The pathology of disease-causing mutations in these genes follow this pattern, with SYT2 dysfunction resulting in muscle weakness without cognitive effects and SYT1 dysfunction sparing NMJs but causing cognitive and behavioral problems. In the current study we characterized synaptic transmission across a range of SYT1 and SYT2 disease-associated alleles using the *Drosophila* model. The most severe disease alleles mapped to the SYT1 C2B Ca^2+^-binding pocket that mediates insertion of its hydrophobic loops into the PM. Although there is a range of phenotypes observed in these patients depending on the specific residue that is mutated, all cause severe intellectual and behavioral phenotypes associated with little to no language acquisition. When the corresponding mutations are expressed in *Drosophila* containing one endogenous wildtype SYT1 allele, they caused a dominant-negative phenotype that reduced evoked neurotransmitter release. Similar effects were observed in SYT2 C2B Ca^2+^-binding pocket mutations. These observations are consistent with prior studies proposing a dominant-negative function for this class of disease alleles^7,32,33,35,36,38,40,41,44,60^. We propose these mutations engage SNAREs at the SV-PM interface during vesicle priming similar to wildtype SYT1. After clamping SNAREs in a prefusion state, they are unable to undergo normal Ca^2+^-dependent PM insertion to trigger fusion, resulting in a dominant disruption to SV release even in the presence of a wildtype SYT1 or SYT2 allele (Figure 8I). Computational modeling supports this hypothesis, identifying C2B Ca^2+^ coordination and membrane insertion defects in these SYT1 and SYT2 alleles. In addition, a prior screen performed in the lab identified intragenic second site suppressor mutations in SYT1 that rescued lethality when C2B Ca^2+^-binding pocket mutations were expressed pan-neuronally in *Drosophila*^7^. These second site mutations often disrupted SYT1-SNARE interactions or Ca^2+^-independent PM binding by the C2B domain, indicating the C2B Ca^2+^-binding pocket alleles are no longer toxic if the mutant protein is not properly positioned at the SV-PM fusion interface. We did not observe defects in spontaneous release rates following overexpression of most of these alleles. Given SYT1’s role in SV endocytosis does not require SNARE binding, it is unlikely alterations in spontaneous release or SV endocytosis play major roles in pathology. In addition, defects in SV docking are unlikely to contribute to the phenotype, as mutations in the C2B Ca^2+^binding pocket expressed in the *Drosophila Syt1* null background can rescue the reduced SV density observed by EM^18^ and restore the readily releasable SV pool assayed with hypertonic stimulation^20^. Rather, our data indicate these alleles disrupt evoked release after SV docking by “clogging” fusion sites through their ability to engage and lock SNAREs in a pre-fusion state.

In contrast to these severe dominant-negative alleles, SYT1 mutations outside of the C2B Ca^2+^-binding pocket acted through distinct mechanisms. Our data indicate the L159R and T196K mutant alleles in the C2A domain act through haploinsufficiency, as both proteins are unstable and degraded when expressed in *Drosophila*. The L159R protein was completely undetectable, while the T196K allele was partially degraded. Mammalian L159R SYT1 also displayed reduced expression when transfected into mouse hippocampal cultures^36^, consistent with our observations. In addition, expression of T196K could not rescue the *Drosophila Syt1* null phenotype. Indeed, we found that *Syt1* null heterozygotes with only one functional SYT1 allele caused a 40% reduction in evoked synaptic transmission compared to controls with two wildtype alleles. Patients with the L159R and T196K alleles display a milder set of phenotypes compared to the SYT1 C2B Ca^2+^-binding pocket patients^35^. The L159R patient was able to use language with good verbal comprehension and was diagnosed with autism spectrum disorder. The T196K patient developed language and had no motor delay, but was diagnosed as hyperactive with occasional self-injury. The M303K SYT1 C2B disease allele also reduces SYT1 expression and may fall into the same loss-of-function category, as this patient displays milder symptoms than other C2B Ca^2+^-binding variants^34^. SYT1 haploinsufficiency as a disease-causing mechanism is supported by recent studies identifying *de novo* chromosomal rearrangements disrupting SYT1 in two independent patients^61,62^. One patient was diagnosed with autism, intellectual disability and speech impairment, while the second had a more complex phenotypic presentation due to the second chromosomal breakpoint causing split-hand foot malformation (SHFM1) disorder. Although our data indicate synaptic transmission is dosage-sensitive for SYT1, haploinsufficiency for SYT2 is unlikely to cause neuromuscular disease in humans. Four rare cases of homozygous recessive loss of SYT2 have been described^63–65,66^, with patients showing profound motor dysfunction. Parents heterozygous for these null alleles did not display any reported neuromuscular defects, indicating 50% of SYT2 levels are sufficient for NMJ function. Given both SYT1 and SYT2 are present on SVs at motor terminals^4^, the loss of one allele of SYT2 appears to be less deleterious than the loss of one copy of SYT1 at CNS synapses where it is often the sole isoform expressed.

A third class of alleles we identified was represented by the C2A E219Q and C2B N341S variants that acted in a gain-of-function manner. The E219Q patient displayed milder phenotypes than the C2B Ca^2+^ pocket mutations, highlighted by ataxic gate, autism and obsessive-compulsive disorder. Together with the T196K allele, the N341S variant resulted in the mildest phenotypes described for a SYT1 variant^35^. When E219Q and N341S were reintroduced into the *Drosophila Syt1* null background, they displayed a robust 3 to 5-fold increase in evoked responses compared to rescue with wildtype SYT1. The N341S allele also caused a dramatic increase in mini frequency. N341 resides within the primary SYT1-SNARE interface^51^ and may alter SNARE interactions. N341 and the nearby Y339 residue form direct interactions with D166 and H162 in one SNAP-25 SNARE helix^58^. As part of this interaction, N341 forms H-bonds with both Y339 and D166. The N341S variant is likely to disrupt the normal H-bond with Y339, allowing Y339 to form stronger stacking interactions with the aromatic ring of H162 in SNAP-25 and potentially strengthening SYT1-SNARE attachments that enhance both evoked and spontaneous SV fusogenicity. A recent study of N341S in rodent neurons also discovered a large increase in spontaneous release caused by this allele, with evidence the serine substitution may be phosphorylated and alter SNAP-25 interactions through this alternative mechanism^67^. Although N341S enhances spontaneous release frequency, we hypothesize the larger effect on evoked release is the main driver of disease pathology. How E219Q mechanistically enhances evoked release is less clear. This negatively charged residue resides at the base of the C2A domain away from the Ca^2+^ binding loops and may counteract the attachment of the C2A domain to the PM in a docked pre-fusion state. SYT1 C2A attachment to the PM normally requires the R234 and K237 basic residues in the C2A Ca^2+^ binding loops and the polybasic K190-K193 stretch on the C2A domain surface^58^. E219 resides near the polybasic stretch that anchors the C2A domain to the PM. Neutralization of this negative charge might increase SYT1 mobility to allow more robust Ca^2+^-induced conformational changes required to drive evoked fusion.

Together, these results reveal a phenotypic spectrum from the most severe alleles that dominantly disrupt synaptic transmission, to weaker haplo-insufficient alleles with milder decreases in evoked release, and gain-of-function alleles that enhance synaptic transmission and cause mild behavioral phenotypes (Figure 8I). Our data indicate C2B Ca^2+^ binding pocket alleles for SYT1 and SYT2 dominantly decrease evoked synaptic transmission by binding to SNARE complexes and the PM while preventing Ca^2+^-dependent membrane insertion that block release sites. Alleles outside of this region can destabilize SYT1 and act via a haplo-insufficient mechanism to reduce evoked release, or hyperactivate SYT1 to generate gain-of-function phenotypes that enhance SV fusion. By defining the underlying mechanisms for each allele, more refined approaches can now be considered for future treatment options. Therapeutics that enhance neurotransmission would be indicated for the SYT1 loss-of-function alleles. In contrast, gain-of-function alleles are more likely to require therapeutics that reduce evoked release to bring synaptic transmission back to a more normal range. The severe category of C2B Ca^2+^ binding pocket variants will likely be the most difficult to treat given the strong-dominant negative effects they generate that block SV fusion and clog release sites. Directed knockdown of the dominant-negative allele will likely be required. If knockdown could be globally achieved and maintained early enough in development, symptoms might be improved to a milder range similar to those observed in patients with SYT1 haploinsufficiency.

## Limitation of the study

Given our analysis was performed using the *Drosophila* model, additional defects in SYT1 function might be present in humans with these variants. How these alterations in synaptic transmission alter brain circuitry during childhood is also unclear but will further compound the initial defects caused by SYT1 dysfunction. We only analyzed how SYT1 and SYT2 mutations affect glutamatergic synaptic transmission, so additional defects might be present at inhibitory synapses in the case of SYT1 or at cholinergic NMJs for SYT2. Given the strong conservation of the SV fusion machinery, one would expect similar defects across most synapse types. How the SYT1 variants alter dense core vesicle release from neuromodulatory neurons is unknown, and defects in this form of neuronal communication could contribute to certain phenotypes in patients as well. Finally, the effects of the more severe C2B Ca^2+^ binding pocket variants on synaptic transmission is likely to be underestimated in our study, as our overexpression system does not generate a full 1-to-1 match of endogenous to mutant SYT1. As such, we would expect more severe defects in evoked release is the protein expression of each allele were equally matched.

## Acknowledgements

We thank the Bloomington *Drosophila* Stock Center (Indiana University, Bloomington, IN; NIH P40OD018537), the Developmental Studies Hybridoma Bank (University of Iowa, Iowa City, IA), and members of the Littleton lab for helpful discussions. This work was supported by the Freedom Together Foundation and a National Institutes of Health grant (NS40296) to J.T.L. Anton2 computer time was provided by the Pittsburgh Supercomputing Center (PSC) by a National Institutes of Health grant (GM116961). The Anton2 machine at PSC was generously made available by D.E. Shaw Research. Stampede and Anvil supercomputer time was provided through allocation from the ACCESS program, which is supported by National Science Foundation grants #2138259, #2138286, #2138307, #2137603, and #2138296.

## Author contributions

Conceptualization, Z.G. and J.T.L.; methodology, Z.G., M.B. and J.T.L., investigation, Z.G. and M.B.; funding acquisition, J.T.L.; supervision, J.T.L.; writing—original draft, Z.G., M.B. and J.T.L.; writing—review & editing, Z.G., M.B., and J.T.L.

---

**Supplemental Figure 1.**
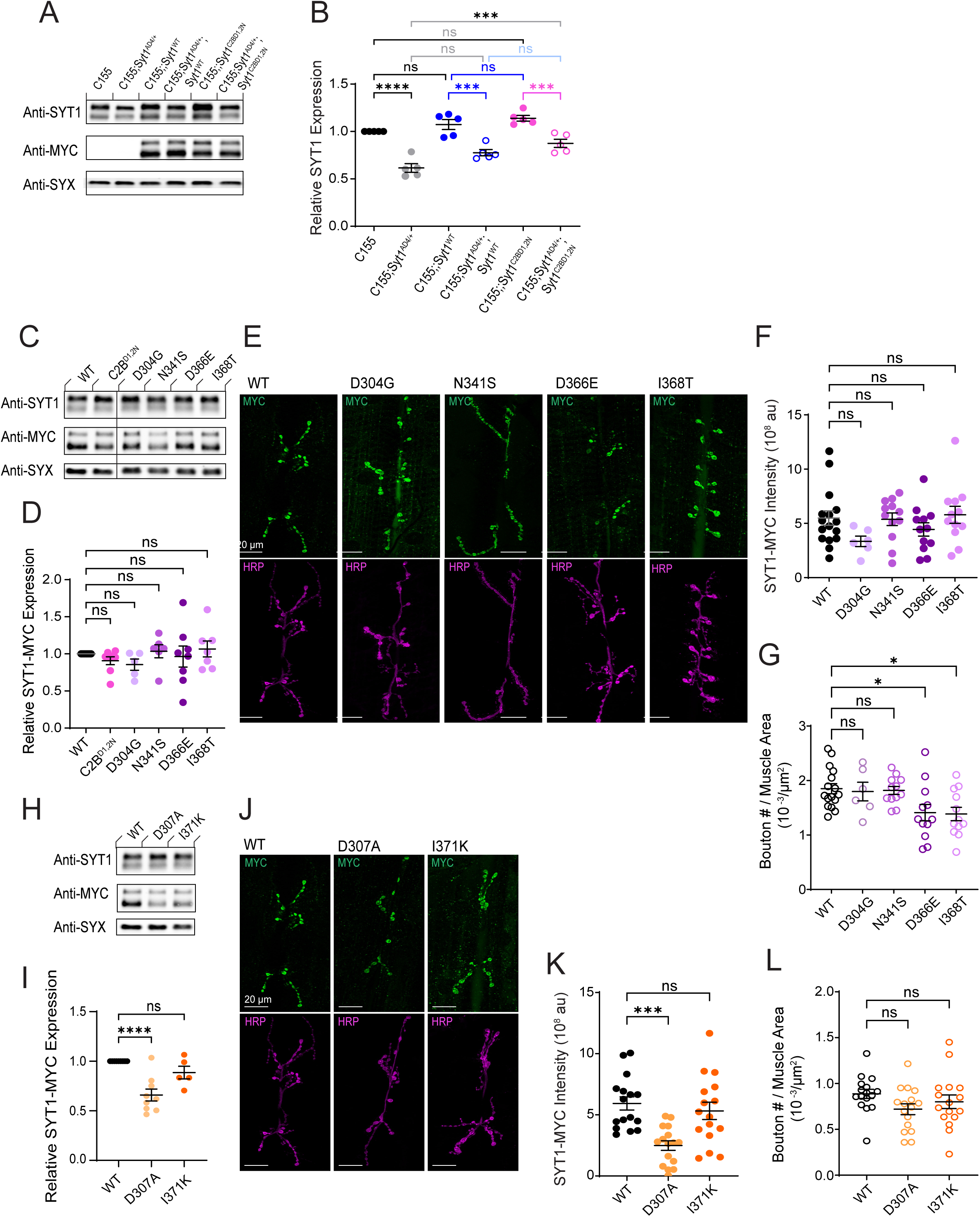
Expression and localization of C2B Ca^2+^ binding pocket SYT1 and SYT2 mutants. (A) Representative western for SYT1 (anti-SYT1) and SYT1-MYC expression (anti-MYC) from adult brain extracts with anti-SYX as a loading control for the indicated genotypes: *C155*, *C155;Syt1^AD4/+^*, *C155;;Syt1^WT^*, *C155;Syt^AD4/+^;Syt1^WT^*, *C155;;Syt1^C2BD1,2N^*, *C155;Syt1^AD4/+^;Syt1^C2BD1,2N^*). (B) Quantification of SYT1 protein expression normalized to control (C155) for the indicated genotypes. (C) Representative western for SYT1 (anti-SYT1) and SYT1-MYC expression (anti-MYC) from adult brain extracts with anti-SYX as a loading control for SYT1 disease alleles: *C155;Syt1^AD4/+^;Syt1^WT^*(WT)*, C155;Syt1^AD4/+^;Syt1^C2BD1,2N^* (C2B^D1,2N^), *C155;Syt1^AD4/+^;Syt1^D304G^* (D304G), *C155;Syt1^AD4/+^; Syt1^N341S^* (N341S), *C155;Syt1^AD4/+^;Syt1^D366E^*(D366E), and *C155;Syt1^AD4/+^;Syt1^I368T^* (I368T). (D) Quantification of SYT1-MYC protein expression normalized to control (WT) for the indicated genotypes. Note the SYT1 C2B disease alleles are overexpressed similarly to the wildtype SYT1 protein. (E) Immunostaining for SYT1 (anti-MYC, green) and HRP (magenta) at 3^rd^ instar muscle 6/7 NMJs of the indicated genotypes from C. (F) Quantification of total SYT1-MYC fluorescence within the HRP-positive NMJ area (au = arbitrary units). Note the SYT1 C2B disease alleles traffic normally to presynaptic terminals (G) Quantification of varicosity number normalized to muscle surface area for the indicated genotypes. (H) Representative western for SYT1 (anti-SYT1) and SYT1-MYC expression (anti-MYC) from adult brain extracts with anti-SYX as a loading control for SYT2 disease alleles: *C155;Syt1^AD4/+^;Syt1^WT^*(WT), *C155;Syt1^AD4/+^;Syt1^D307A^* (D307A) and *C155;Syt1^AD4/+^;Syt1^I371K^*(I371K). (I) Quantification of SYT1-MYC protein expression normalized to control (WT) for the indicated genotypes. (J) Immunostaining for SYT1 (anti-MYC, green) and HRP (magenta) at 3^rd^ instar muscle 6/7 NMJs of the indicated genotypes from panel I. (K) Quantification of total SYT1-MYC fluorescence within the HRP-positive NMJ area (au = arbitrary units). (L) Quantification of varicosity number normalized to muscle surface area for the indicated genotypes. Data are shown as mean ± SEM. Statistical significance: * *p* < 0.05, *** *p* < 0.001, **** *p* < 0.0001, ns = not significant. Raw values, statistical tests and sample number are provided in the Source Data file.

**Supplemental Figure 2.**
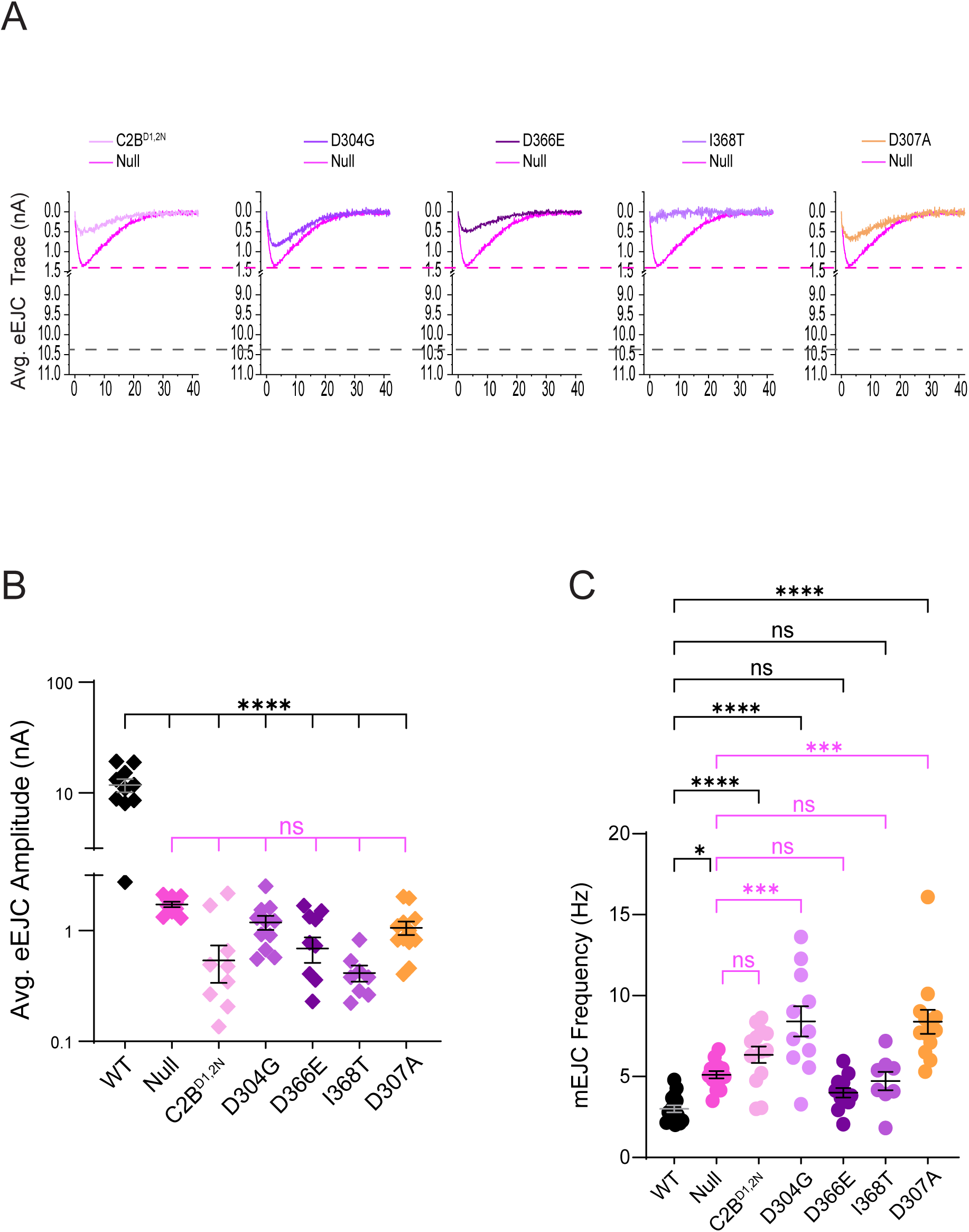
Disease-causing mutations in the C2B Ca^2+^ binding loops fail to rescue *Syt1* null phenotypes. (A) Average eEJC traces for the indicated genotypes: *C155;Syt1^AD4/N13^;Syt1^C2B-D1,2N^* (C2B^D1,2N^), *C155;Syt1^AD4/N13^;Syt1^D304G^* (D304G), *C155;Syt1^AD4/N13^;Syt1^D366E^* (D366E), *C155;Syt1^AD4/N13^;Syt1^I368T^*(I368T) and *C155;Syt1^AD4/N13^;Syt1^D307A^* (D307A). The average mean EJC trace for the *Syt1* null mutant is shown in magenta with its mean EJC amplitude represented by the dashed magenta line. The mean EJC amplitude for rescue of the null mutant with WT SYT1 is represented by the dashed grey line. (B) Quantification of mean eEJC amplitudes for the indicated genotypes. Note the C2B disease alleles fail to rescue evoked release defects in the *Syt1* null background. (C) Quantification of mean spontaneous release rate for the indicated genotypes. Data are shown as mean ± SEM. Statistical significance: * *p* < 0.05, *** *p* < 0.001, **** *p* < 0.0001, ns = not significant. Raw values, statistical tests and sample number are provided in the Source Data file.

**Supplemental Figure 3.**
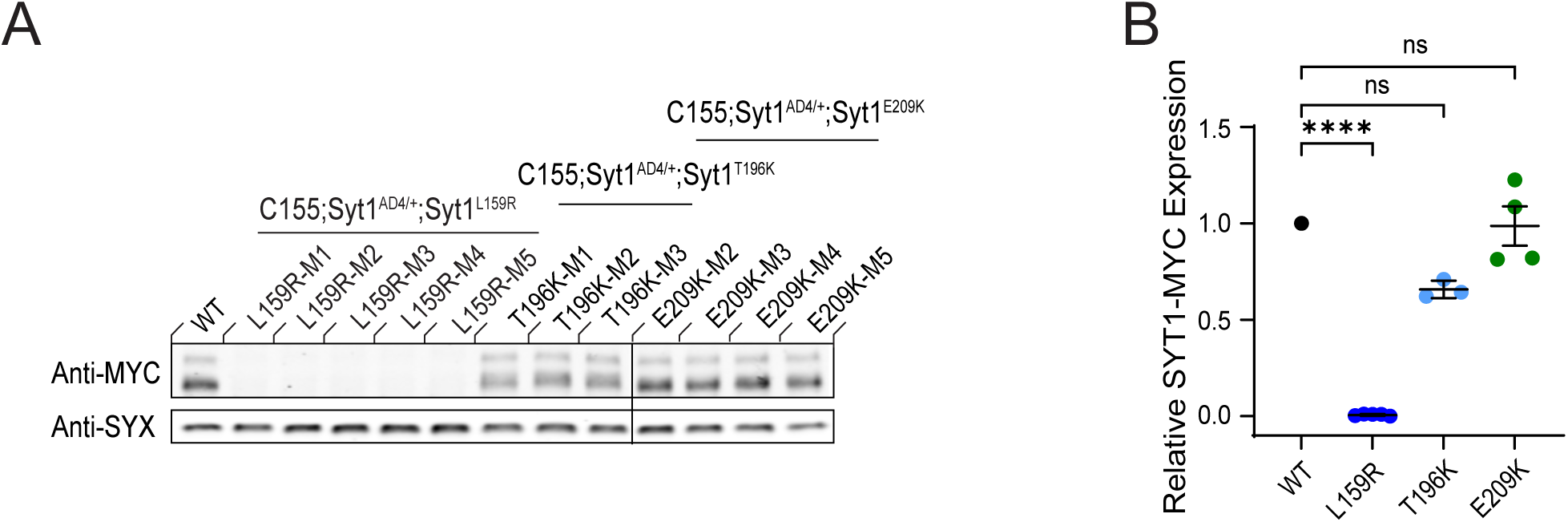
Reduced SYT1 protein levels for the C2A L159R and T196K disease alleles. (A) Western for SYT1-MYC expression (anti-MYC) from adult brain extracts with anti-SYX as a loading control for independent transgenic lines of the indicated genotypes: *C155;Syt1^AD4/+^;Syt1^WT^* (WT), *C155;Syt1^AD4/+^;Syt1^L159R^*(L159R), *C155;Syt1^AD4/+^;Syt1^T196K^* (T196K), and *C155;Syt1^AD4/+^;Syt1^E209K^* (E209K). (B) Quantification of SYT1-MYC protein expression normalized to control (WT) for the indicated genotypes. Each data point represents an independent transgenic line run in a separate lane from the blot. Note the dramatic reduction in protein levels for the L159R allele. Data are shown as mean ± SEM. Statistical significance: **** *p* < 0.0001, ns = not significant. Raw values, statistical tests and sample number are provided in the Source Data file.

## STAR Methods

### KEY RESOURCES TABLE

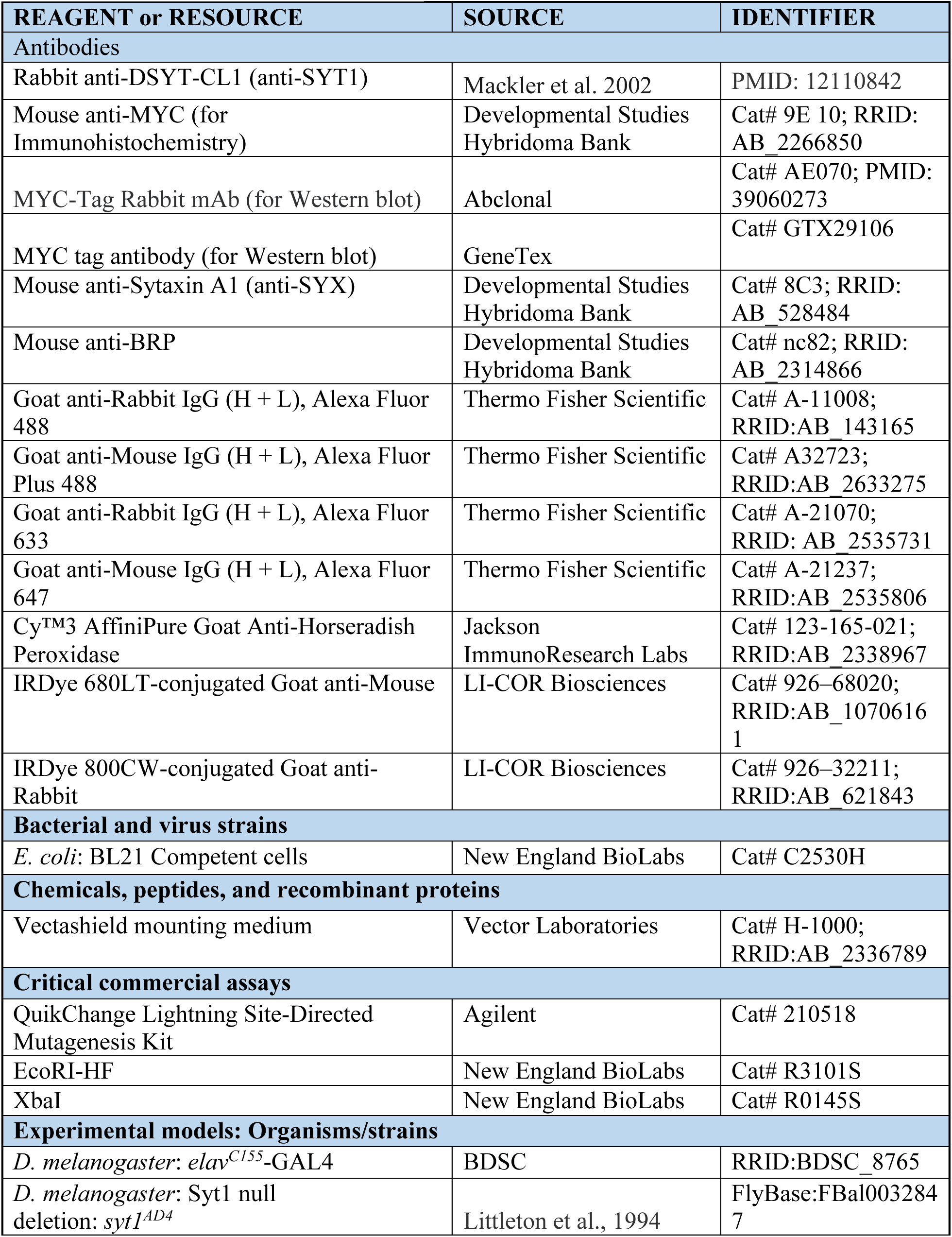
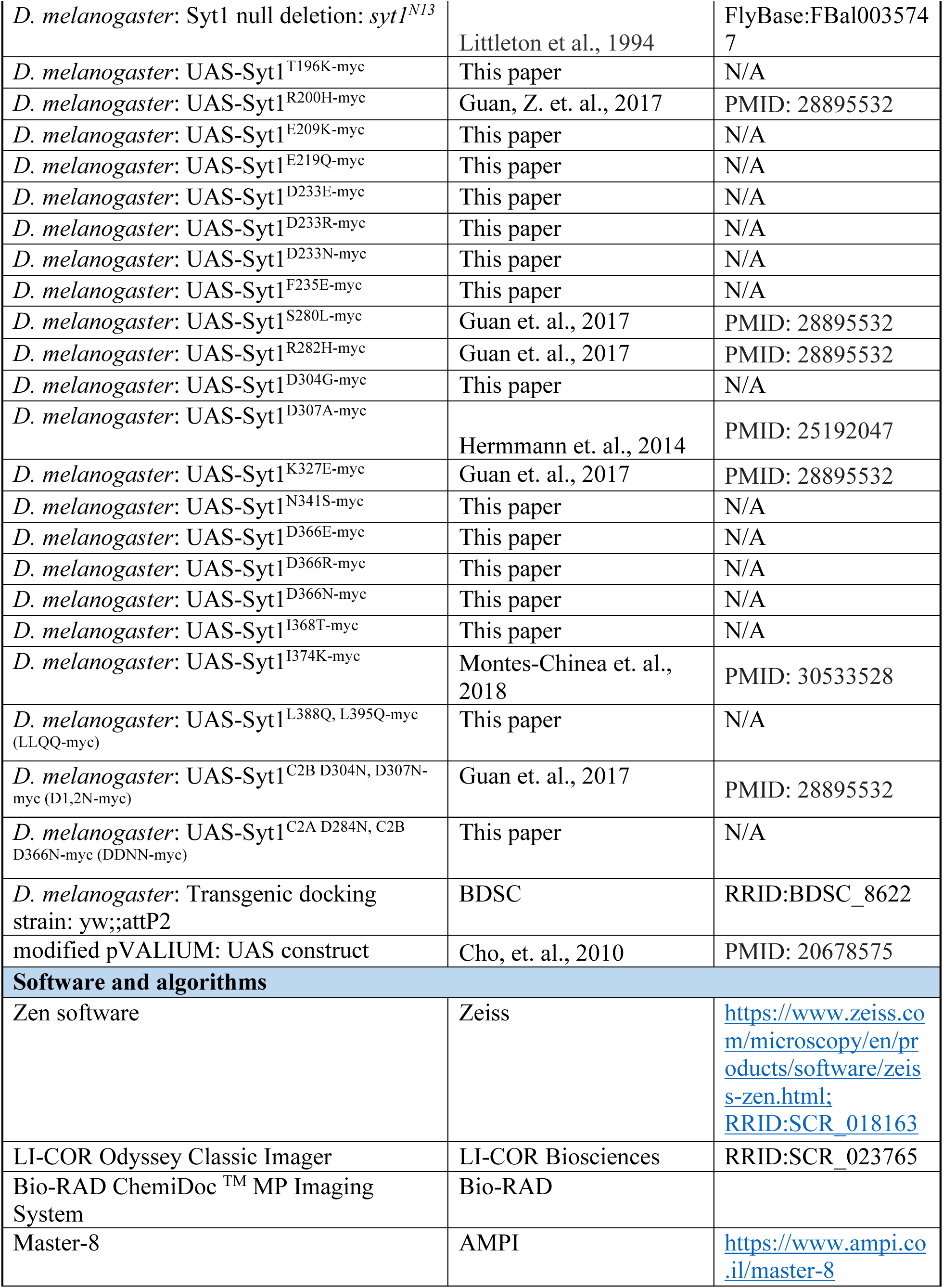
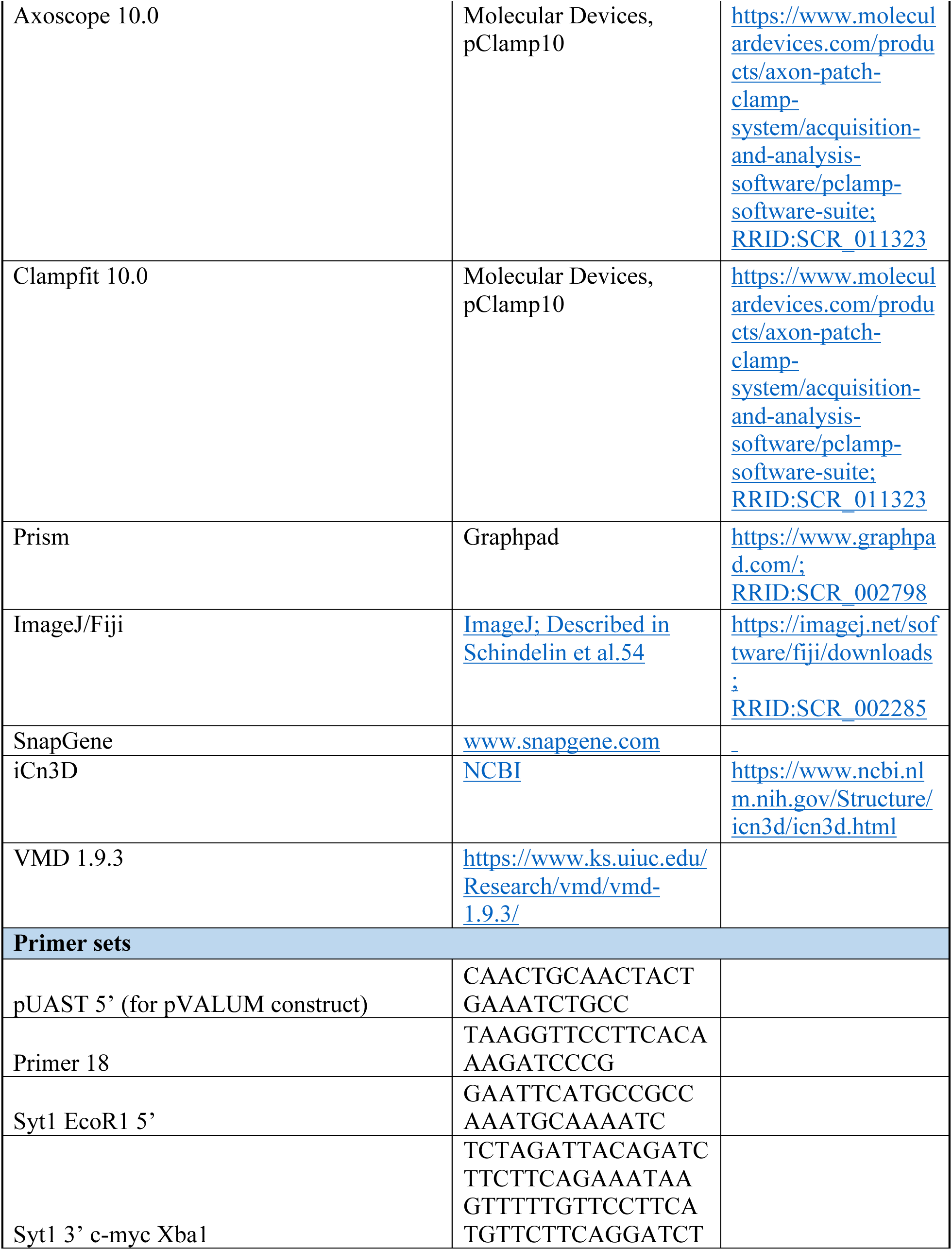
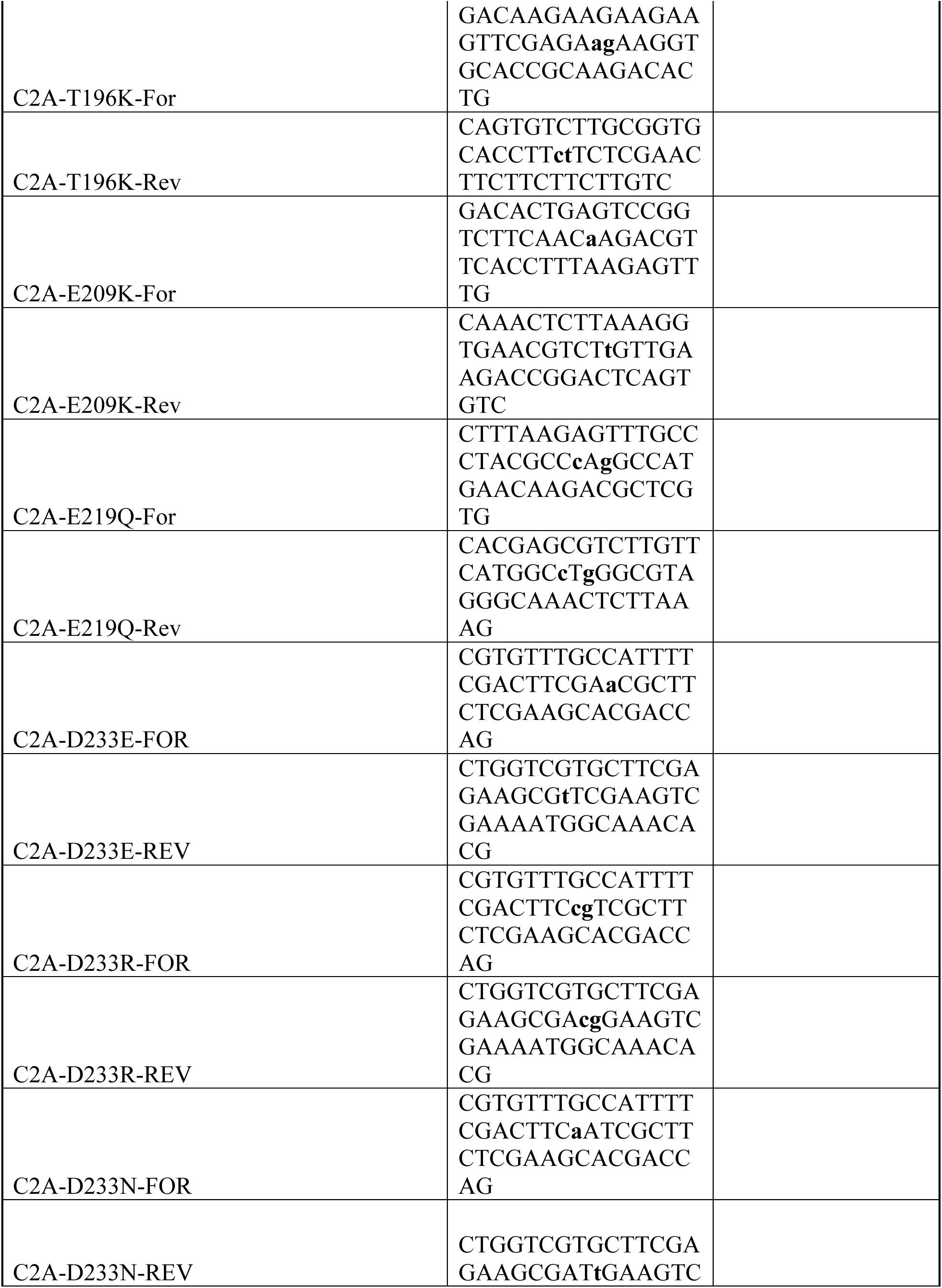
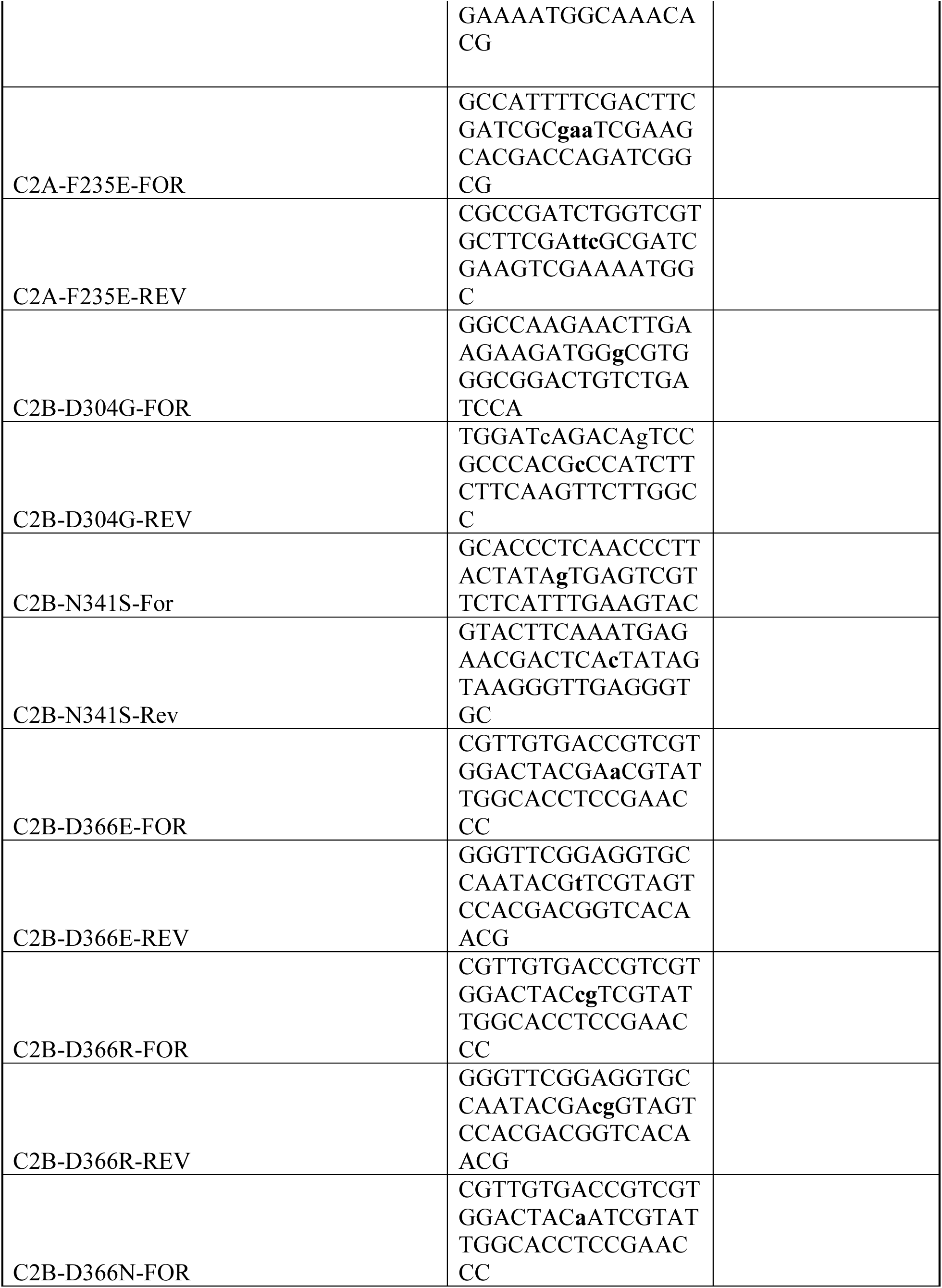
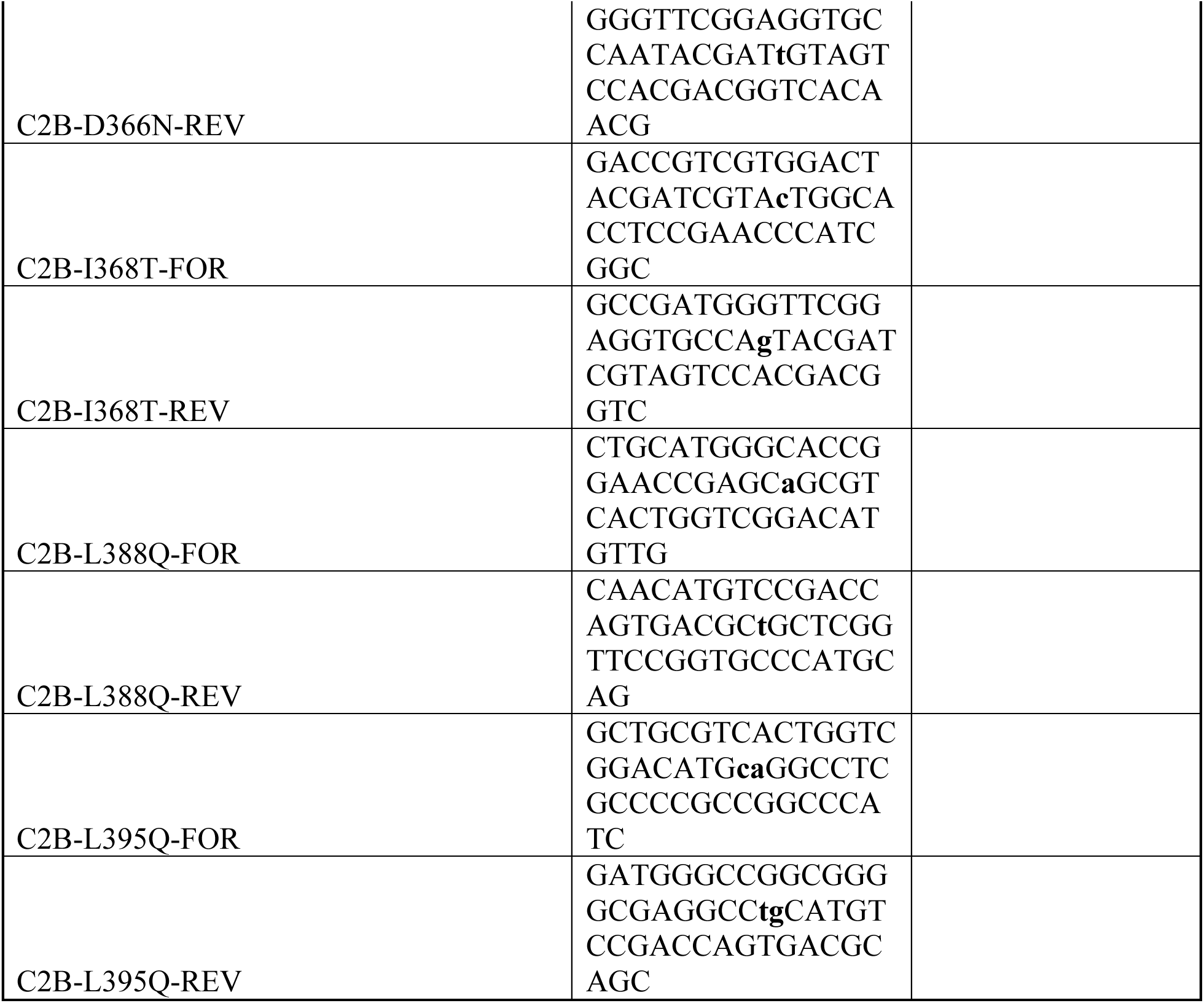

### RESOURCE AVAILABILITY

#### Lead contact

Further information and requests for resources and reagents should be directed to and will be fulfilled by the lead contact, J. Troy Littleton.

#### Materials availability

*Drosophila* stocks generated in this study are available from the lead contact upon request without restriction.

#### Data and code availability

- This paper does not report any original code.
- Any additional information required to reanalyze the data reported in this paper is available from the lead contact upon request.

### EXPERIMENTAL MODEL AND STUDY PARTICIPANT DETAILS

#### Animals

*Drosophila melanogaster* were cultured on standard medium and maintained at 25°C. Late 3^rd^ instar larvae in the wandering stage were used for imaging and electrophysiological experiments. Western blots were performed on adult brain extracts from animals aged 3 days. Males were used for experiments unless otherwise noted. Experiments were performed in a *w^11^*^18^ (BDSC #3605) genetic background unless otherwise noted. Genotypes of the strains used are reported in the figure legends and indicated in the resource table. For electrophysiology, experimenters were blinded to genotype for both data collection and analysis. No vertebrate animals were used in this study, so institutional permission and oversight, health/immune status, participants involvement in prior procedures, and drug or naive state, are not applicable.

### METHOD DETAILS

#### Transgenic constructs

QuikChange Lightning Site-Directed Mutagenesis Kit (Agilent) was used for site-directed mutagenesis on pVALUM-SYT1-MYC to generate *Syt1* mutations tagged with MYC at the C-terminus. These were subcloned into the modified pVALIUM construct and injected into a yw;;attP2 third chromosome docking strain by BestGene Inc (BDSC #8622). UAS lines were recombined into the *Syt1^AD4^* null mutant heterozygous background. *Syt^AD4/N^*^13^ was used as the *Syt1 null* mutant background, and *elav^C1^*^55^-GAL4 (BDSC #8765) was used for pan-neuronal transgene expression.

#### Immunohistochemistry

Larvae were dissected in hemolymph-like HL3.1 solution (in mM: 70 NaCl, 5 KCl, 4 MgCl_2_, 10 NaHCO3, 5 trehalose, 115 sucrose, 5 HEPES, pH 7.2) and fixed in 4% paraformaldehyde for 20 min. Larvae were washed three times for 5 min with PBST (PBS containing 0.1% Triton X-100), followed by a 1 hour incubation in block solution (5% NGS (normal goat serum) in PBST). Fresh block solution and primary antibodies were then added. Samples were incubated overnight at 4°C and washed with two short washes and six extended 120 min washed in PBST. PBST was replaced with block solution and fluorophore-conjugated secondary antibodies were added. Samples were incubated overnight at 4°C. Finally, larvae were rewashed with PBST and mounted in Vectashield (Vector Laboratories). Antibodies used for this study include: rabbit anti-SYT1 1:1000 (provided by Noreen Reist); mouse anti-MYC for Immunohistochemistry (1:200, Developmental Studies Hybridoma Bank (DSHB) Cat# 9E 10); mouse anti-SYX1 (1:1000, DSHB Cat# 8C3); mouse anti-BRP, 1:500 (DSHB Cat# NC82); Cy™3 AffiniPure Goat Anti-Horseradish Peroxidase (HRP) (1:1000, Jackson ImmunoResearch Labs, Cat# 123-165-021). Fluorophore-conjugated secondary antibodies were used at 1:400.

#### Confocal imaging and imaging data analysis

Imaging was performed on a Zeiss Pascal confocal microscope (Carl Zeiss Microscopy) using a 63×1.3 NA oil-immersion objective (Carl Zeiss Microscopy). Images were processed with the Zen (Zeiss) software. A 3D image stack was acquired for each NMJ imaged (muscle 6/7 NMJ of abdominal segment A3) and merged into a single plane for 2D analysis using Volocity (x64) image analysis software. No more than two NMJs were analyzed per larva. Anti-HRP labeling was used to identify neuronal anatomy (axons and NMJs) and quantify synaptic bouton number and NMJ area. Muscle 6/7 NMJ area was used to normalize quantifications for muscle surface area. For SYT1-MYC fluorescence quantification, the HRP-positive area was used to outline NMJs and axons. Total SYT1-MYC fluorescent intensity was measured in the outlined area, with background fluorescence of mean pixel intensity of non-HRP areas subtracted.

#### Two-electrode voltage-clamp electrophysiology

Postsynaptic currents were recorded from 3^rd^ instar muscle 6 at segment A3 using two-electrode voltage clamp with a −80 mV holding potential. Experiments were performed at room temperature HL3.1 saline solution (in mM, 70 NaCl, 5 KCl, 10 NaHCO3, 4 MgCl2, 5 trehalose, 115 sucrose, 5 HEPES, pH 7.2). Final [Ca^2+^] was adjusted to 0.5 mM unless otherwise noted. Motor axon bundles were cut and suctioned into a glass electrode and action potentials were stimulated at 0.5 Hz using a programmable stimulator (Master-8, AMPI). Data acquisition and analysis was performed using Axoscope 10.0 and Clampfit 10.0 software (Molecular Devices) and inward currents were labeled on a reverse axis for clarity. Graphs were performed using Origin Software (OriginLab Corporation, Northampton, MA). Average EJC traces were generated in Clampfit 10 using the Analyze -> Average Traces. The resulting average traces were copied starting from the 0 nA baseline and imported into Origin 2020. In Origin 2020, traces from all samples within the same experimental group were aligned by time and averaged to generate the group average EJC trace.

#### Western blot analysis

Westerns were performed using standard laboratory procedures with rabbit anti-SYT1 (1:1000) or rabbit anti-MYC (1:1000, Abclonal Cat# AE070; or GeneTex Cat# GTX29106), incubating with mouse anti-SYX1 (1:1000, DSHB Cat# 8C3). IRDye 680LT-conjugated goat anti-mouse (1:5000, LI-COR Biosciences 926–68020) and IRDye 800CW-conjugated goat anti-rabbit (1:5000, LI-COR Biosciences 926–32211) were used as secondary antibodies. Blocking was performed with Blocking Buffer (Rockland Immunochemicals) overnight at 4°C. Primary antibodies were diluted in the Blocking Buffer and incubated overnight at 4°C. Secondary antibodies were diluted in a solution containing four parts TBST (1X TBS with 1% Tween 20) to one part Blocking Buffer incubating for overnight at 4°C. A LI-COR Odyssey Imaging System (LI-COR Biosciences) or Bio-RAD ChemiDoc TM MP Imaging System was used for visualization. Analysis was performed using FIJI image analysis software. Relative SYT1-MYC expression was calculated by normalizing to SYX1 intensity.

#### Molecular dynamics

Molecular systems were constructed using Visual Molecular Dynamics Software (VMD, Theoretical and Computational Biophysics Group, NIH Center for Macromolecular Modeling and Bioinformatics, at the Beckman Institute, University of Illinois at Urbana-Champaign). Simulations were performed in a water/ion environment with explicit waters. Potassium and chloride ions were added to neutralize the systems and to yield 150 mM concentration of KCl. Water boxes with added ions were constructed using VMD. The initial structure of the anionic lipid bilayer containing phosphatidylserine (POPS) and PIP_2_, POPC:POPS:PIP_2_ (75:20:5)^68^ was kindly provided by Dr. J. Wereszczynski (Illinois Institute of Technology) and positioned in the XY plain. The initial structure for the C2B domain^69^ in its Ca^2+^-bound form was obtained from crystallography studies (Protein Data Bank #1TJX). The initial structure of the Ca^2+^ free protein was obtained by removing Ca^2+^ ions. The protein was positioned near the lipid bilayer at a random orientation such that it did not interact with lipids. The molecular system included 83265 atoms with a periodic cell size of 82x86x116 Å. Single point mutations were introduced using VMD. The initial configurations were identical for all C2B variants. To generate Ca^2+^-bound C2B modules, when two K^+^ ions were transiently trapped inside of the Ca^2+^-binding pocket at the end of the trajectory, K^+^ ions were replaced by Ca^2+^ and the system was stripped of K^+^ and Cl^-^, re-ionized, and re-equilibrated^59^.

MD simulations were performed employing CHARMM36 force field^70^ modified to include the parameters for PIP_2_ as described. The simulations were performed with periodic boundary conditions and Ewald electrostatics in NPT ensemble at 310K. Heating (20 ps) and equilibration (50 ns) phases were performed employing NAMD^71^ Scalable Molecular Dynamics (Theoretical and Computational Biophysics Group, NIH Center for Macromolecular Modeling and Bioinformatics, at the Beckman Institute, University of Illinois at Urbana-Champaign) at ACCESS (Advanced Cyberinfrastructure Coordination Ecosystem: Services & Support) Stampede and Anvil clusters. NAMD simulations were performed with a flexible cell and a time-step of 1.5 fs, employing Langevin thermostat and Berendsen barostat. Production runs were performed at Anton2 supercomputer^72,73^ with Desmond software through the MMBioS (National Center for Multiscale Modeling of Biological Systems, Pittsburg Supercomputing Center and D.E. Shaw Research Institute). All Anton2 simulations were performed in a semi-isotropic regime, with a time-step of 2.5 fs, and employing the multigrator^74^ to maintain constant temperature and pressure. The trajectory analysis was performed with VMD. All parameters along the trajectories were computed with a time step of 2.4 ns.

#### Replication of results

Western blot experiments were carried out in at least four biological replicates containing three adult fly brains per sample. For confocal imaging and TEVC recordings, no more than two NMJs were analyzed per larva. All genetic crosses were set up at least twice to obtain reproducible results from replicate to replicate.

### QUANTIFICATION AND STATISTICAL ANALYSIS

#### Experimental design and statistical analysis

Statistical analysis and plot generation was performed using GraphPad Prism software version 9.5.1. Appropriate sample size was determined using a normality test. Statistical significance for comparisons of two groups was determined by an unpaired t-test. For comparisons of three or more groups of data, one-way analysis of variance (ANOVA) (nonparametric) with post hoc Sidak multiple comparisons test. The *p* values associated with 1-way ANOVA tests were adjusted *p* values obtained from a post hoc Sidak multiple comparisons test. Appropriate sample size was determined using GraphPad Statmate. In all figures, the data are presented as mean ± SEM. Statistical comparisons are with control, unless noted. The results were all shown: ns = no significant change (*p* > 0.05), * *p* < 0.05, ** *p* < 0.005, *** *p* < 0.001, and **** *p* < 0.0001. All error bars are SEM.

